# Site-specific adaptation mechanisms of an oral pathobiont in the oral and gut mucosae

**DOI:** 10.1101/2022.01.13.476151

**Authors:** Yijie Guo, Sho Kitamoto, Gustavo Caballero-Flores, Daisuke Watanabe, Kohei Sugihara, Gabriel Núñez, Christopher J. Alteri, Naohiro Inohara, Nobuhiko Kamada

## Abstract

Periodontal inflammation leads to oral dysbiosis with the expansion of oral pathobionts. Besides the pathogenic role of oral pathobionts during periodontal inflammation, studies have revealed that oral pathobionts contribute to diseases in distant organs beyond the oral mucosa. For example, the oral pathobiont *Klebsiella aerogenes*, which accumulates in the oral mucosa during periodontitis in mice, can exacerbate colitis when it ectopically colonizes the gastrointestinal tract. However, the precise mechanisms by which oral pathobionts establish their colonization in extra-oral mucosal sites remains incompletely understood. We performed high-throughput *in vivo* genetic screening to identify fitness genes required for the adaptation of the oral pathobiont *K. aerogenes* to different mucosal sites – the oral and gut mucosae – at the steady state and during inflammation. In addition, the global transcriptome of *K. aerogenes* in different environments was analyzed. We determined that *K. aerogenes* employs genes related to iron acquisition and chaperone usher pili, which are encoded on a newly identified genomic locus named “locus of colonization in the inflamed gut” (LIG), for adaptation in the gut mucosa, particularly during inflammation. In contrast, the LIG virulence factors are not required for *K. aerogenes* to adapt to the oral mucosa. Thus, oral pathobionts likely exploit distinct adaptation mechanisms in their ectopically colonized intestinal niche compared to their original niche.

**Author Summary:** The symbiotic bacterial community evolves uniquely at each body site. However, under certain circumstances, symbionts may disseminate far from their original niche and colonize a site ectopically. It has been reported that oral resident bacteria associated with periodontal disease, namely oral pathobionts, contribute to gastrointestinal diseases, such as inflammatory bowel disease and colorectal cancer, through ectopic colonization of the gut. However, the molecular mechanisms by which oral pathobionts adapt to the gut environment remain largely unexplored. In this study, using a model oral pathobiont *Klebsiella aerogenes*, we screened genes essential for the colonization of the pathobiont at different mucosal sites. We discovered that the oral pathobiont uses distinct virulence mechanisms at two different mucosal sites – the oral and gut mucosae. Understanding the strategies of bacteria localized in ectopic organs may lead to the development of strategies to prevent disease caused by the dissemination of pathogenic oral bacteria.

## Introduction

The oral cavity is a complex environment that nourishes a large number of microorganisms that colonize the surfaces of teeth and the mucosal tissue to maintain oral and systemic health [1]. It houses the second most diverse and abundant microbial community, next to the gastrointestinal (GI) tract, harboring over 770 species of bacteria [2]. Oral microbes play a vital role in homeostasis and disease. The symbiotic oral microbiota, which can enhance oral mucosal immunity, displays colonization resistance that prevents the colonization by invasive species that cause oral disease [3, 4]. In contrast, certain oral bacteria, such as *Porphyromonas gingivalis*, *Tannerella forsythia*, *Fusobacterium nucleatum*, and *Aggregatibacter actinomycetemcomitans*, and species of *Treponema* and *Prevotella*, are enriched in patients with oral disease and participate in the pathogenesis of the disease [5–7].

Accumulating evidence has confirmed that certain oral bacteria can ectopically colonize the GI tract and cause disease, such as inflammatory bowel disease (IBD) and colorectal cancer [8, 9]. It has been reported that IBD patients display an increased prevalence of periodontitis [10, 11]. Also, early periodontitis may worsen clinical outcomes in some IBD patients [12]. Furthermore, the oral administration of *P. gingivalis* and *F. nucleatum* exacerbates the severity of dextran sulfate sodium (DSS)–induced colitis in mice by disrupting the intestinal barrier [13–15]. Investigators have observed that *Enterobacteriaceae* (especially *Klebsiella* spp.) isolated from the saliva of IBD patients can ectopically colonize the gut, inducing pathogenic Th1 cell differentiation and contributing to gut disease [16]. In our research, we have found that oral inflammation promotes the ectopic gut colonization of oral bacteria. In a mouse periodontitis model, *Klebsiella* and *Enterobacter* spp. accumulated with oral dysbiosis and translocated to the lower digestive tract. The ectopic colonization of oral-derived *Klebsiella* and *Enterobacter* spp. increased the susceptibility to DSS-induced colitis [17]. Therefore, oral pathobionts can colonize and promote inflammation in distant organs, such as the GI tract. Given the difference in the microenvironment between the oral and gut mucosae, as well as differences in the diversity and abundance of competing microbes, oral pathobionts need to use a different strategy from their natural colonization in the oral mucosa to establish their ectopic colonization in the gut mucosa. However, the mechanisms by which oral pathobionts adapt to the gut environment remain largely unclear.

Here, to identify bacterial genes that facilitate the oral pathobionts ability to successfully colonize and survive in the different mucosal sites, we used random transposon mutagenesis combined with high-throughput sequencing techniques to simultaneously screen thousands of insertion mutants for fitness defects in two different microenvironments between the oral and gut mucosae during steady state and inflammation [18–20]. In this context, we used *K. aerogenes* as a model oral pathobionts, as we and others reported *Klebsiella* spp. reside in the oral cavity in humans and mice, particularly during periodontal disease [16, 17, 21, 22]. More importantly, oral *Klebsiella* spp. contribute to the pathogenesis of IBD through their ectopic gut colonization in both humans and mice [16, 17]. Using the oral pathobiont *Klebsiella aerogenes* SK431, we created a saturating library of transposon mutants and conducted transposon–insertion site sequencing (Tn-Seq) in murine models of periodontitis and colitis caused by or exacerbated by the colonization of the oral pathobiont. Using Tn-Seq, we discovered various virulence genes, including those for iron acquisition, fimbrial adhesins, chaperon usher pili, and the type 6 secretion system (T6SS), all associated with the colonization of *K. aerogenes* SK431 in the oral and gut mucosae. Moreover, we identified a newly genomic locus in *K. aerogenes* required for adaptation to the inflamed gut mucosa. Thus, oral pathobionts appear to exploit distinct adaptation strategies in their original niche compared to their ectopically colonized niche – the gut.

## Materials and Methods

### Bacteria and Mice

The oral pathobiont *K. aerogenes* SK431 [17] was cultured in Luria Broth (LB) medium and stored at -80 °C in 25 (v/v) % glycerol until use. Specific pathogen-free (SPF) and germ-free (GF) C57BL/6 (B6) mice (age 8–12 weeks, female and male) were obtained from The Jackson Laboratory (Bar Harbor, Maine) and bred under SPF and GF conditions in our mouse facility at the University of Michigan. All SPF mice were regularly maintained on conventional rodent chow (Laboratory Rodent Diet 5001, LabDiet, St. Louis, MO). The gut microbiotas of SPF mice were normalized by exchanging their bedding 2-3 times a week for 2 weeks before use. All GF mice were housed in flexible film isolators, provided with distilled water ad libitum, and fed Autoclavable Rodent Breeder Diet 5013 (LabDiet). Their GF status was checked weekly by aerobic and anaerobic culture. The absence of the microbiota was verified by microscopic analysis of stained cecal contents, which detects any unculturable contamination. All animal study protocols were approved by the Institutional Animal Care & Use Committee at the University of Michigan.

### Transposon mutant library generation and inoculation

The transposon mutant library was generated by the conjugative transfer of the suicide plasmid pSAM_Cam, carrying a mariner-based transposon flanking the kanamycin resistant (Kan^R^) cassette [18, 19], from the donor strain *E. coli* S17-1 (Kan^R^, ampicillin sensitive [Amp^S^]) to the recipient *Klebsiella aerogenes* strain SK431 (Ampicillin resistant [Amp^R^], Kanamycin sensitive [Kan^S^]). A library of random transposon mutants was created by mating a mid-log phase culture of *E. coli* S17-1 with a pre-heat shock (20 min, 42 °C) mid-log phase culture of *K. aerogenes* SK431 at a 1:1 ratio. Mating mixtures were centrifuged (4,500 rpm, 10 min), washed with PBS, spread onto 0.22 μm filter discs (MilliporeSigma, Burlington, MA), plated on LB agar plates with 250 μM IPTG (Thermo Fisher Scientific, Waltham, MA), and incubated at 37 °C for 5 h. Then, the filter discs were transferred to tubes and rinsed with 1 mL PBS. The bacterial suspensions were plated onto LB agar plus 50 µg/mL Kan and 100 µg/mL Amp and incubated at 37 °C overnight to isolate the SK431 mutants. The mutants were flooded with PBS using a cell spreader. Multiple conjugation experiments were conducted, and mutants from multiple plates were harvested and pooled. The pooled stock solution, contains 63,000 SK431 mutant colonies, was stored in 25 % glycerol at -80 °C until needed. To prepare the inoculum (i.e., input), an aliquot of the frozen stock was thawed at 4 °C and culture in LB medium at 37 °C with shaking until the culture reached an OD_600_ 0.6 – 0.8.

### Experimental periodontitis model in mice

SPF and GF B6 mice were used for Tn-Seq and RNA-Seq analyses, respectively. Periodontitis was induced by ligature placement [23]. Briefly, silk suture thread (SUT-15-1; Roboz Surgical Instrument Co., Gaithersburg, MD) was soaked with the 10^9^ CFU *K. aerogenes* SK431 WT strain or SK431 Tn mutant library in PBS for 3 h, and then inserted between the first and second maxillary molars on the contralateral right and left sides (i.e., ligature placement). The oral ligatures remained for 1 day (control group) or 14 days (inflamed group). Mice with periodontitis (14 day-ligature) or without periodontitis (1-day ligature) were euthanized, and the oral ligatures were removed. In the Tn-Seq experiment, the ligatures were suspended in PBS, plated onto LB agar plus 50 µg/mL Kan and 100 µg/mL Amp, and incubated at 37 °C overnight to harvest SK431 mutants colonized in the oral cavity (i.e., oral ligatures). Bacterial colonies were flooded with PBS using a cell spreader for bacterial DNA extraction. In the RNA-Seq experiment, ligature-associated bacteria were collected and immediately used for RNA extraction. Maxilla tissues were used for histological evaluation of periodontitis. Tissue biopsies from the maxillae were fixed in 4% paraformaldehyde. Alveolar bone loss was measured by micro–computed tomography (micro-CT) and visualized by MicroView Standard software (Parallax Innovations, 2.5.0–3139), and sections were stained with hematoxylin and eosin (HE) for histological analysis. Histological scores were assigned blindly by a trained pathologist who graded the severity of periodontitis from 0 to 4 as described [23–26].

### Experimental colitis model in mice

A dextran sulfate sodium (DSS)–induced colitis model was used in mice. SPF and GF B6 mice were used for Tn-Seq and RNA-Seq analyses, respectively. For the Tn-Seq analysis, colitis was induced in SPF mice by administering 2 (w/v) % DSS for 6 days. On day 6 post DSS treatment, the SK431 Tn mutant library (1 × 10^9^ CFU/mouse) was administered by oral gavage. Colonized SK431 mutant strains were then harvested 3 days post challenge. Harvested feces were plated on LB agar plus 50 µg/mL Kan and 100 µg/mL Amp and incubated at 37 °C overnight to isolate SK431 mutants. Mutants were flooded with PBS using a cell spreader and used for bacterial DNA extraction. For the RNA-Seq analysis, GF B6 mice were colonized with SK431 WT strain (1 × 10^9^ CFU/mouse) for 2 weeks and then challenged with 1.5% DSS for 5 days. On day 5 post DSS challenge, feces were collected for bacterial RNA extraction. Isolated colonic tissue was fixed in 4% paraformaldehyde, and sections were stained with HE. Histological scores were assigned blindly by a trained pathologist who evaluated two variables [27]: severity of inflammation (0–16) and extent of epithelial/crypt damage (0–16). An overall colitis score (0–32) was obtained by summing the score assigned to each variable.

### Genome analysis of *K. aerogenes* SK431

DNA was isolated from the overnight culture of SK431 and subjected to random sequencing by Illumina MiSeq as described [28]. The resulting paired-end sequences were assembled to contigs by SPAdes [29]. The genes of SK431 and other reference genomes, whose complete sequences were obtained from NCBI GenBank, were annotated by Prokka [30]. Putative virulence genes were predicted by BLASTP with virulence factor database (VFDB) entries [31]. Orthologue gene groups and the phylogenetic tree distances were determined by roary [32]. The Newick file of the resulting phylogenetic tree was visualized by TreeGraph 2 [33]. To compare Tn mutation and mRNA expression frequencies at pathway level among groups, total TnPKM and FPKM of pathways were calculated based on the subsystem categories of RAST [34]. Synteny maps were obtained using SyntTax [35].

### Mutant library sequencing and bioinformatic analyses

Bacterial DNA was purified from the pathogen library grown *in vitro* (input), and mutants recovered from selected LB plates after collection from the ligature (oral) and fecal (gut) samples (output) using the DNeasy Blood & Tissue Kit (Qiagen Sciences, Germantown, MD) [36]. Preparation of the libraries for Illumina sequencing was performed following a published protocol [37]. Briefly, the NEB Ultra II FS kit (NEB #E6177L, New England Biolabs, Ipswich, MA) was used for enzymatic fragmentation of genomic DNA, end-repair of fragments, and ligation of the adaptor. Then, the transposon–genome junctions were enriched by PCR and indexed using NEBNext Multiplex Oligos (NEB #E7600S, New England Biolabs). The libraries were quantified by the University of Michigan Advanced Genomics Core using the Agilent TapeStation system (Agilent, Santa Clara, CA) and by quantitative PCR (qPCR) using a KAPA Library Quantification Kit for Illumina sequencing platforms (Kapa Biosystems, Roche, Wilmington, MA). Samples were multiplexed and sequenced with a 15% spike-in of PhiX DNA in an Illumina NovaSeq platform (Illumina, San Diego, CA) using a 200 cycle single-end read at the same facility. Tn sequences were mapped to the genome of SK431 [17] using Bowtie 2 [38] and annotated/counted by HTSeq) [39]. Required genes were defined as the relative Tn abundance per kilobase gene length per million reads (TnPKM) in the output divided by the input. Total frequencies of the pathways were calculated based on the subsystem categories of RAST [34] using signal-to-noise ratios (SNR) between two groups as determined by Morpheus (https://software.broadinstitute.org/morpheus/).

### Bacterial transcriptome analysis

Bacterial RNA samples isolated from the ligature (oral) and fecal (gut) samples were used for microbial RNA-seq. Total RNA was extracted using a phenol:chloroform:isoamyl alcohol mixture (pH 4.5; AM9720, Invitrogen, Thermo Fisher Scientific, Waltham, MA) and LiCl precipitation solution (final concentration to 2.5 M, Invitrogen). The isolated RNA was treated with DNase (Invitrogen) and RNeasy (Qiagen Sciences). Any host RNA and bacterial ribosomal RNA were depleted using the NEBNext rRNA Depletion Kits (New England BioLabs). Isolated RNA was then assessed for quality using the Agilent TapeStation system. 5 ng of total RNA was used for intact RNA library preparation (NEBNext Ultra II Directional RNA Library Prep Kit for Illumina (NEB #E7760, New England BioLabs). To obtain sufficient RNA for the analysis of oral samples, we pooled RNA samples purified from two mice. The mRNA was fragmented and converted into cDNA using reverse transcriptase and random primers [40]. The products were purified and enriched by PCR to create the final cDNA library. Final libraries were checked for quality and quantity using the Agilent TapeStation system and qPCR using a KAPA Library Quantification Kit for Illumina sequencing platforms (Kapa Biosystems, Roche). Sequencing was conducted in the Illumina NovaSeq platform (Illumina) using 150 cycle pair-end reads at the University of Michigan Advanced Genomics Core. Adapters and low-quality bases were removed from the raw data. The Pair-end reads were mapped to the SK431 genomic sequence using Bowtie 2 [38] with default parameters after the removal of reads mapped to the Illumina control phiX174 and to reference genomes (Genome ID: 548.747). Then, reads of individual genes were counted by HTSeq [39]. Gene expressions were shown using fragments per kilobase transcript per million reads (FPKM). Total frequencies of the pathways were calculated based on the subsystem categories of RAST [34] using SNR between two groups as determined by Morpheus .

### Quantification and statistical analysis

Statistical analyses were performed using GraphPad Prism 8 software (GraphPad Software, San Diego, CA). Differences between two groups were evaluated using a 2-sided Student *t* test and Mann–Whitney *U* test for parametric and non-parametric datasets (determined by Shapiro-Wilk test and *F* test), respectively. Statistical significance in highly repetitive comparison between two groups was evaluated by false positive discovery rate (FDR) analysis using Morpheus. Non-metric multidimensional scaling (NMDS) plots were generated using Bray-Curtis dissimilarity indexes. Differences in Bray-Curtis dissimilarity indexes were evaluated by permutational multivariate analysis of variance (PERMANOVA) using PAST3 [41]. Differences at *P* < 0.05 and FDR < 0.05 were considered significant.

## Results

### Characterization of the *K. aerogenes* SK431 strain

Our previous study confirmed that the oral pathobiont *K. aerogenes* (Ka) strain SK431 in the inflamed oral cavity of periodontitis mice can ectopically colonize the gut and trigger or exacerbate colitis [17]. However, the genetic features of this strain remain unclear. Therefore, we determined genomic sequences of SK431 and compared the genetic characteristics of this strain with the reference Ka strain KCTC 2190 and 17 other Ka strains identified in the NCBI GenBank database (**Fig. 1**) by roary [32]. Ka strain SK431 showed high evolutionary relationships to AR0161, a poorly described Ka strain, among the different Ka strains studied (**Fig. 1A** and **S1 Fig**), as 94 % of SK431 gene orthologues were found in AR0161. The putative virulence genes of Ka strains were predicted by homology in the entries in the VFDB database [31] and compared with related Enterobacteriaceae pathogens (**Fig. 1B**). Ka SK431 and other 18 reference Ka strains have similar sets of putative virulence genes (**Fig. 1B**). All Ka strains have common systems associated with iron acquisition, including Sit (inner membrane-located iron/manganese transporter), TonB-dependent enterobactin (Ent/Fep), salmochelin (Sal), aerobactin (Aer), and heme-iron acquisition (Chu), suggesting conservation of efficient iron acquisition of Ka in host-associated environments [42]. Also, all Ka strains harbor type VI secretion systems (T6SS), but lack type III secretion system (T3SS) and type IV secretion system (T4SS), which are involved in competition with other bacteria and host immune invasion or intoxication, respectively [43]. Ka SK431 also harbors strain-specific competition factors, including a group A colicin (A-Col) and contact-dependent growth inhibition (Cdi) systems. Ka SK431 genome harbors several putative adhesion factors, including orthologues of known pili (Fim, CfA, *Escherichia coli* common pilus [Ecp]), and conserved but uncharacterized chaperon usher (CU) pili systems (named CUP1 to CUP6 here, **Fig. 1B**). Notably, Sit, Chu, and CUP1 operons were located within a single locus of the SK431 genome (**Fig. 1C**). Other Ka strains, including AR0161, also carry this locus in their genomes, whereas the CUP operon is separated from Sit and Chu in other *Klebsiella* species (**Fig. 1C**), suggesting the evolutionary relationship of this putative virulence locus in the genus *Klebsiella*. As this newly identified genetic locus was composed of the genes that are required for the colonization in the inflamed gut (see below), we named it “locus of inflamed gut colonization” (LIG). These findings suggest that Ka SK431 possesses LIG and other virulence genes conserved in Ka strains. These conserved genes may be responsible for the colonization and virulence of this oral pathobiont at specific sites.

**Fig 1.**
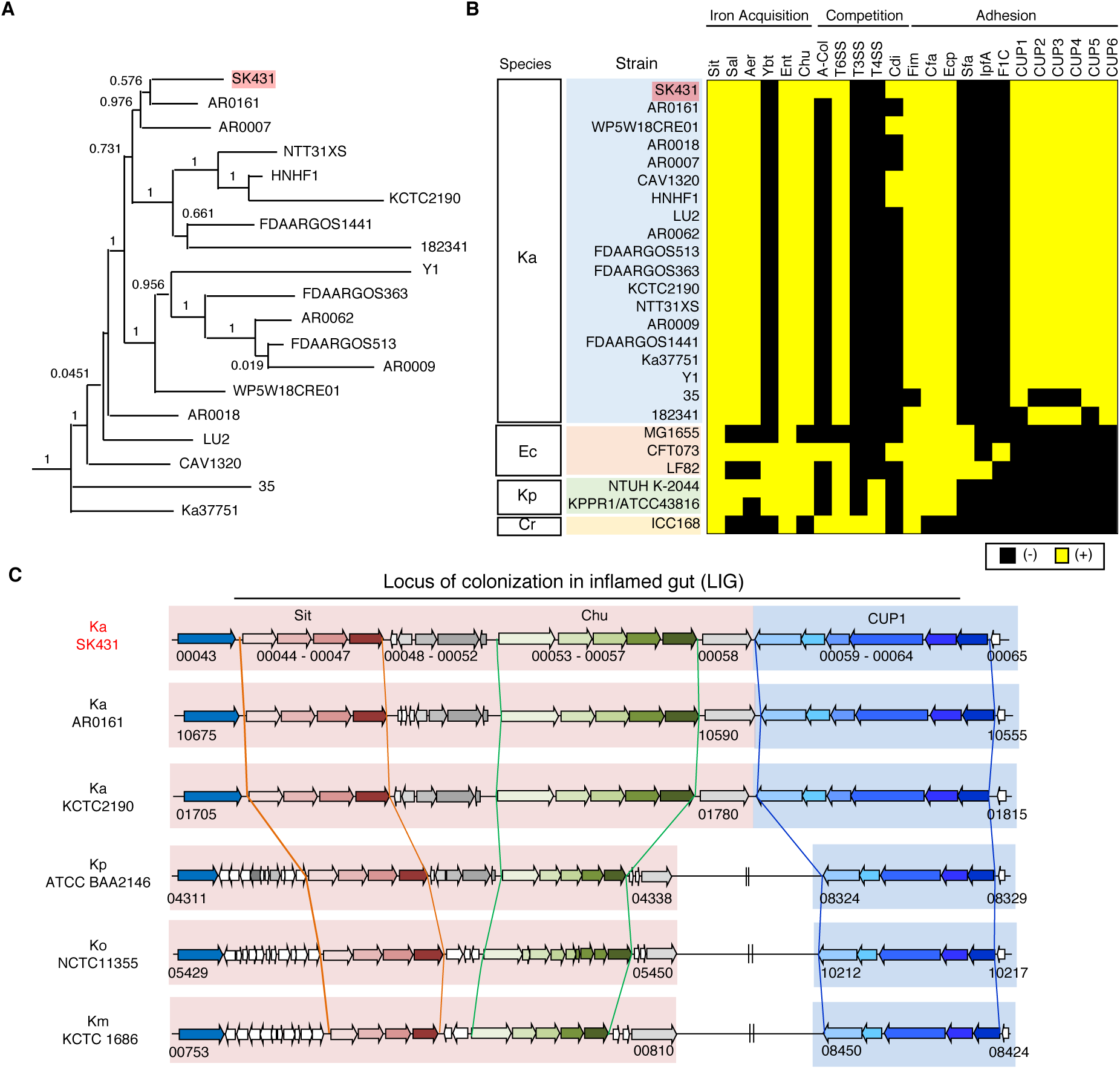
Characterization of *K. aerogenes* SK431 genome. (A) Phylogenetic tree of *Klebsiella aerogenes* SK431 and 18 reference *K. aerogenes* strains. (B) Presence of virulence-associated genes in indicated strains of *Klebsiella aerogenes* (Ka); *Escherichia coli* (Ec); *K. pneumoniae* (Kp); Cr, *Citrobacter rodentium* (Cr). The presence and absence of operon orthologues are indicated by yellow and black, respectively. (C) Schematic representation of “locus colonization in the inflamed gut” (LIG) virulence factors in SK431 and other *Klebsiella* strains. Syntenic regions and gene orthologues are indicated by the same arrows and background colors. The genes of uncharacterized proteins are shown in white arrows. Direction of arrows indicates the transcriptional direction. The locus numbers in gene-annotated genomes (GenBank accession numbers, CP028951, NC015663, 000364385, 900478285, 000240325 for Ka AR0161, Ka KCTC2190, Kp ATCC BA2146, Ko NTCC 11355, Km KCTC1686, respectively) are indicated below the respective first and last genes of each syntenic region.

### Identification of *K. aerogene*s SK431 genes required for the colonization in the oral and gut mucosae

To identify the genes required for the colonization of the oral pathobiont Ka SK431 in its natural (oral) and ectopic (gut) niches, both in the steady state and during inflammation, we generated the SK431 mutant library by insertion of mariner-based transposon (Tn) [44], comprising ∼63,000 independent mutants. To identify genes responsible for the colonization in the healthy and inflamed oral environments, we inoculated the SK431 mutant library with dental ligatures into the oral cavity [17] (**Fig. 2A**). The SK431 mutant strains were then harvested from the inserted ligatures on day 1 (before the development of periodontitis) and day 14 (after the development of periodontitis). To mimic ectopic gut colonization by the oral pathobiont, we also performed orogastric inoculation of the SK431 mutant library (**Fig. 2A**). In the gut environment, the SK431 mutant strains were harvested from feces. DNA was extracted from recovered bacteria (outputs) and the original culture (input; IN) and subjected to Illumina sequencing using the Tn-specific primer (**Fig. 2A**). The outputs include ligature 1-day (control, non-inflamed oral cavity; nOC), ligature 14 days (periodontitis, inflamed oral cavity; iOC), feces of water-treated mice (control, non-inflamed gut; nGT), and feces of DSS-treated mice (colitis, inflamed gut; iGT) samples (**Fig. 2A**). Illumina sequencing analysis showed that the library contained a sufficient number of Tn inserts of individual genes of Ka SK431 and no obvious locus-based bias on a Ka reference genome (**Fig. 2B**). The overall Tn insertion frequencies in particular genes of the outputs (i.e., nOC, iOC, nGT, iGT) were lower than the IN, indicating that these genes were required for optimal colonization at the periodontal site or in the gut (**Fig. 2B** and **S1 Table**). Next, we performed a coverage analysis of the fold difference between two groups to identify genes required to colonize the tested *in vivo* conditions (**Fig. 2C** and **2D**, and results are summarized in **Fig. 2E**). First, we compared the input (IN) and the output samples to identify the genes required for *in vivo* colonization. As shown in **Fig. 2C** and **2E**, numbers of genes are enriched in Group A (IN) compared to Group B (output), indicating that mutants of these genes have reduced fitness in the output groups (i.e., genes are required for *in vivo* adaptation). Notably, the number of required genes was higher in the inflamed gut compared to the input (IN/iGT) than the other conditions (IN/nOC, IN/iOC, and IN/nGT), although the level of requirement was mild to moderate (the majority of genes showed 10- to 25-fold enrichment in IN) (**Fig. 2C** and **2E**). Comparison of the oral and gut environments showed that in both non-inflamed and inflamed conditions, more genes were required for gut colonization than for oral colonization, with mild to moderate requirement levels (the majority of genes showed 10- to 25-fold enrichment in nGT and iGT) (**Fig. 2D** and **2E**). In comparing the non-inflamed and inflamed conditions in each mucosal site (**Fig. 2D** and **2E**), several genes were found to be required for colonization in the inflamed condition than in the steady state for both the oral and gut mucosae. Notably, a small number of genes was highly required (i.e., >100-fold enrichment in nGT) for the colonization in the inflamed gut compared to the non-inflamed gut (iGT vs nGT) (**Fig. 2D** and **2E**). These results suggested that the pathobiont *K. aerogenes* SK431 requires more genes to adapt to the ectopic niche (gut) compared to its natural niche (oral), particularly during inflammation.

**Fig 2.**
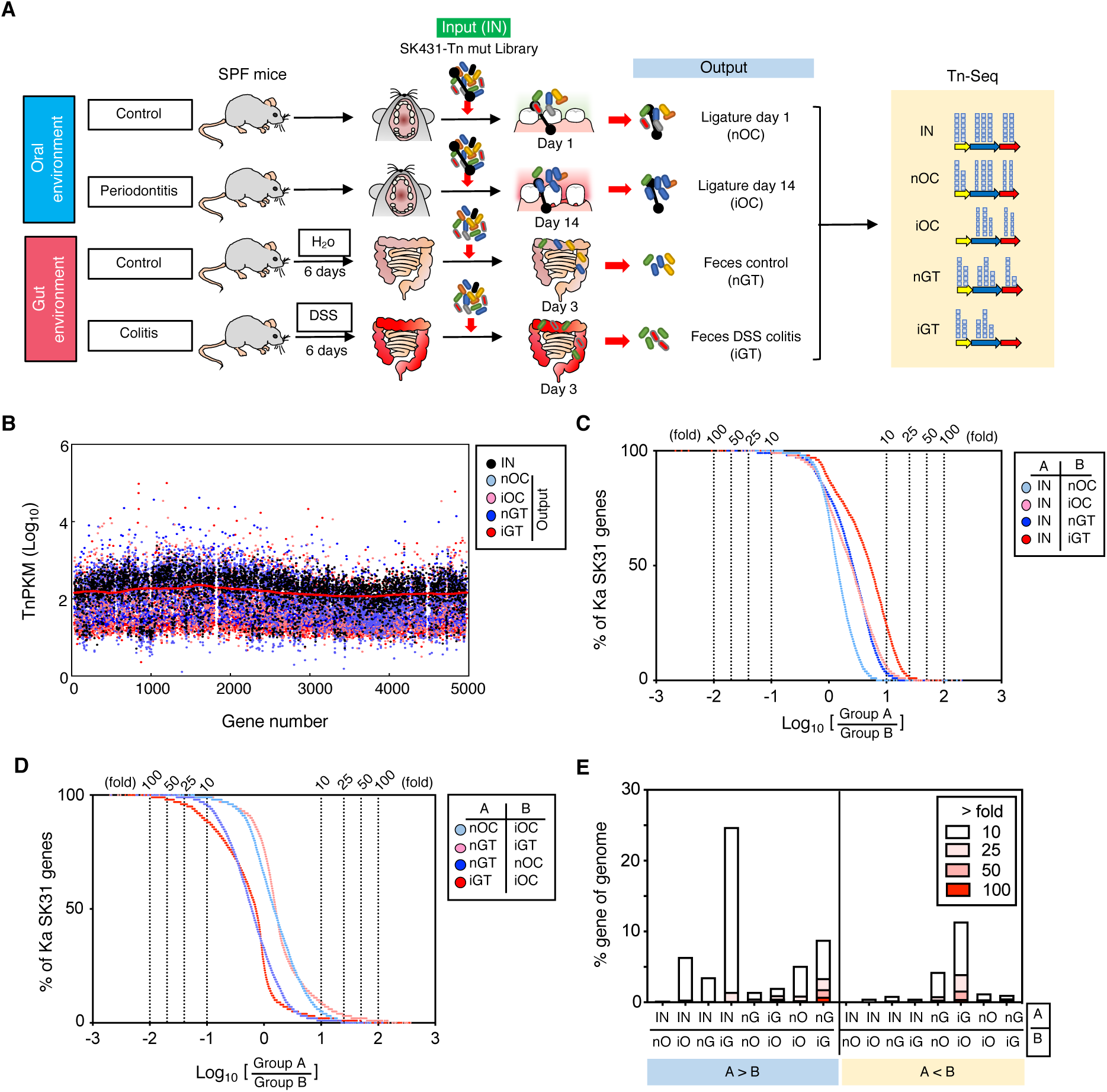
Experimental design and the overview of genes required for the colonization in the steady state and the inflamed oral and gut mucosae. (A) A transposon (Tn) mutant (mut) library of *K. aerogenes* SK431 was inoculated into specific pathogen-free (SPF) B6 mice by soaked ligature and oro-gastric gavage for the oral and gut sites, respectively. Ligature and feces were collected from the healthy control mice and the inflamed (periodontitis or colitis) mice on indicated days after inoculation (output). DNA was isolated from the input and output samples and sequenced by Illumina NovaSeq. The resulting reads were mapped to the Ka SK431 genome, and the abundance of reads at each insertion site from all output samples were compared to those determined for the input samples to determine a fold-change value for each gene. (B) Genome-wide distribution and frequency of mutations in the library. Individual Tn densities per kilobase gene per million reads (TnPKM) are shown as dots on the map of KCTC2190 complete genome, instead of incomplete SK431 genome. Red line, 3,000 gene moving average of IN. (C) (D) The number of genes that were required for the colonization in the indicated locations and conditions. The log ratio of the average TnPKM of indicated Group A per Group B for individual genes is shown on the X-axis. The accumulation curves of the number of the gene from the top (rich in Group B) to the bottom (rich in Group A) are shown on the Y-axis. (E) A graph summarizing the results of (C) and (D).

### Identification of metabolic pathways required for colonization in the oral and gut mucosae

To identify the metabolic pathways that are required for colonization in the healthy and inflamed oral and gut environments, we first obtained the TnPKM sum for the entire metabolic pathways and compared to those of paired groups by SNR (IN vs iOC [**Fig. 3**, open bars] and IN vs iGT [**Fig. 3**, filled bars]). Pathways required for nitrogen metabolism and quinone cofactors synthesis were significantly enriched in the input compared to iOC, suggesting that these pathways are required for colonization of Ka SK431 in the inflamed oral mucosa (**Fig. 3**). Likewise, pathways associated with triacylglycerol metabolism, DNA metabolism, inorganic sulfur assimilation, sulfur metabolism, proline and 4-hydroxyproline metabolism, branched-chain amino acids (AA) metabolism, glycogen metabolism, pyridoxine, capsular and extracellular polysaccharides synthesis, cell wall and capsule synthesis, phosphorus metabolism, iron acquisition and metabolism, and cation transports were significantly enriched in the input compared to iGT (**Fig 4**), indicating the importance of these pathways in the ectopic gut colonization of Ka SK431, especially in the inflamed condition.

**Fig 3.**
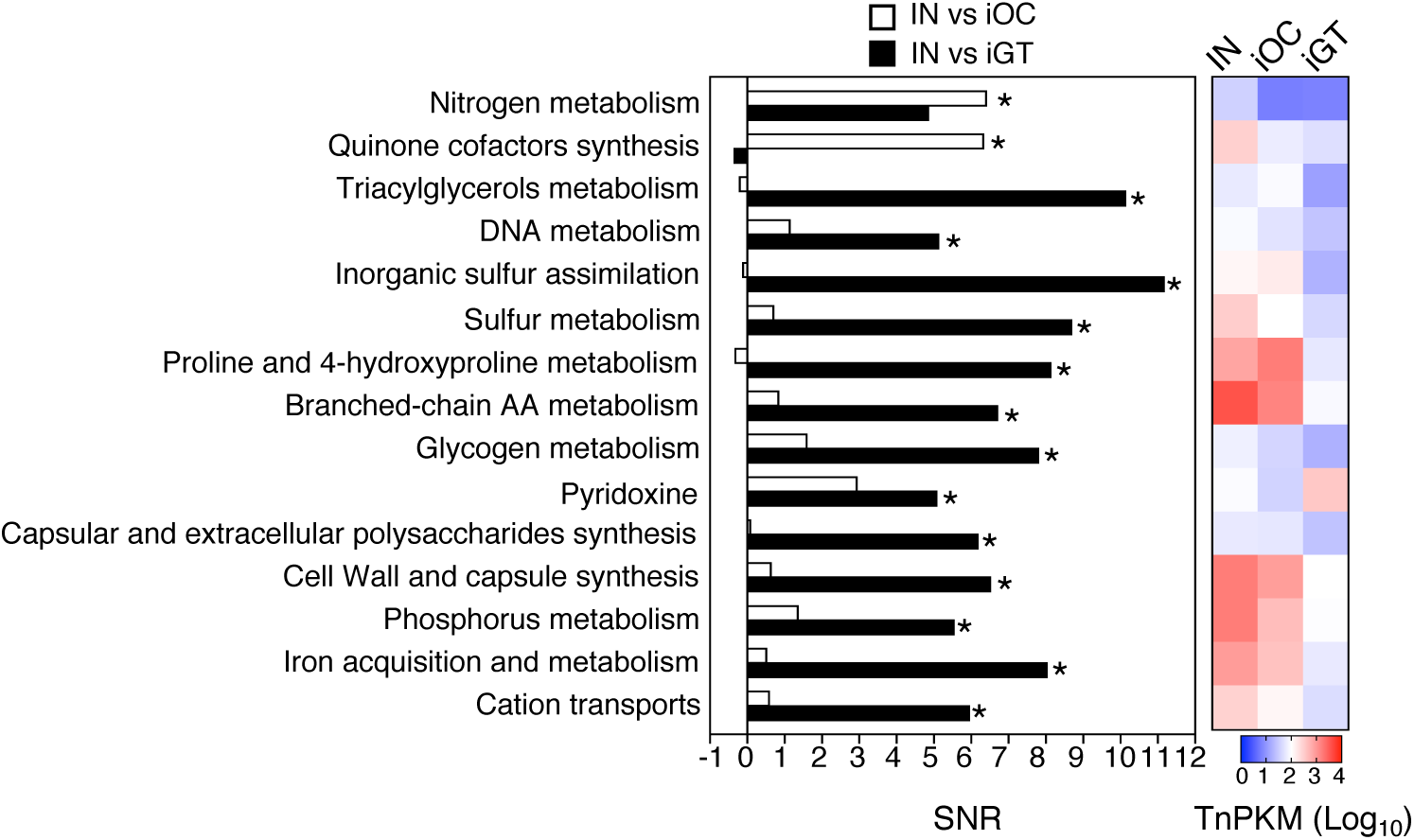
Functional pathways required for the colonization in the oral and gut mucosae. The genes were grouped in functional categories and an SNR analysis was performed using input/output ratios of TnPKM sums (left). The heatmap (right) shows the transposon mutant gene abundance (log_10_ averaged TnPKM, n = 4 to 6 mice) in the input (IN), the inflamed oral cavity (iOC), and the inflamed gut (iGT). The positive values of SNR indicate more in input. *, Statistically significant TnPKM ratio (fold differences > 10, Absolute SNR (|SNR|) > 5, *P* < 0.05, FDR < 0.05). AA, amino acid; TnPKM, transposon (Tn) densities per kilobase per million mapped reads; SNR, signal-to-noise ratio; FDR, false discovery rate.

**Fig 4.**
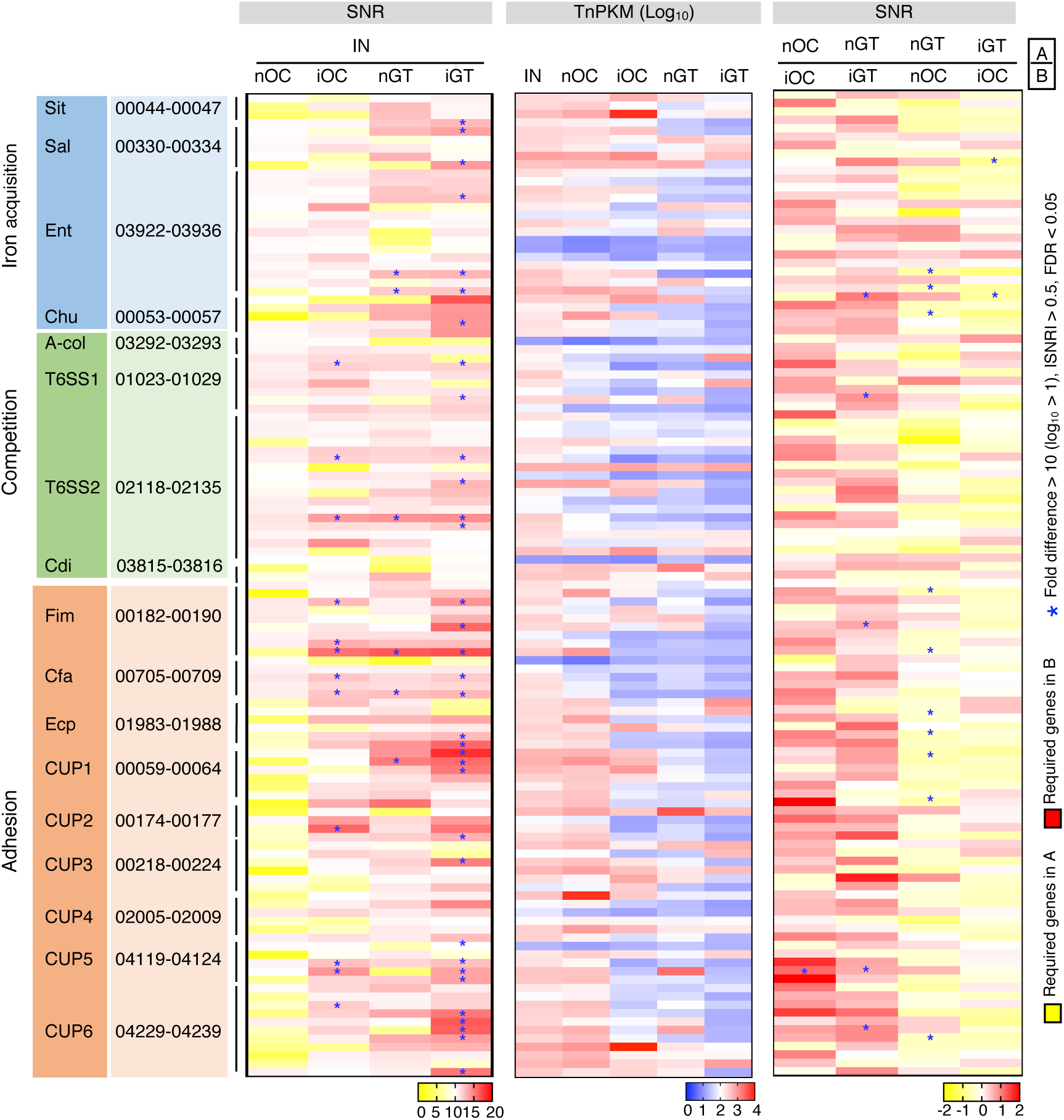
Virulence genes required for SK431 colonization in differential mucosal sites. The statistical difference (SNR) and relative Tn abundance (log_10_TnPKM) of virulence gene mutants of indicated groups were shown by heatmaps (left and right panels for SNR, the middle panel for log_10_TnPKM). The abbreviations of functional systems are shown in Fig.1. The genes ID of each operon were list. T6SS1 and T6SS2, two core operons of T6SS. *, Genes with significant difference (fold differences > 10, |SNR| > 0.5, FDR < 0.05, *P* < 0.05).

### Identification of virulence factors required for the colonization in the oral and gut mucosae

SK431 is an oral pathobiont that can ectopically colonize the gut, causing colitis [17]. Like other bacterial strains belonging to the Enterobacteriaceae family, SK431 harbors various virulence-related factors (**Fig. 1B**). However, it remains uncertain which virulence factors are required to colonize the original niche (i.e., the oral mucosa) and the ectopic niche (i.e., the gut mucosa). To identify virulence genes required for mucosal site-specific adaptation, we analyzed the relative abundance of mutants in virulence genes between the input and outputs. We found that several genes were required to colonize the oral and gut mucosae (**Fig. 4**, left). Several genes were identified as essential for mucosal colonization in the inflamed condition. For example, putative competition factors (T6SS1 and T6SS2) and adhesion factors (Fim, Cfa, CUP2, CUP5, and CUP 6) are required for the colonization in the inflamed oral cavity (iOC) (**Fig. 4**, left). Likewise, genes related to iron acquisition (Sit, Sal, Ent, Chu), competition (T6SS1 and T6SS2), and adhesion (Fim, Cfa, Ecp, CUP1, CUP3, CUP5, and CUP6) were necessary for the colonization in the inflamed gut (iGT). Notably, most of these genes are essential only in the inflamed condition, not in the steady state, in neither the oral nor the gut mucosae, suggesting that SK431 uses virulence factors to adapt to the inflamed mucosal environment. We also attempted to identify the virulence genes required to adapt to the inflamed mucosa by measuring SNRs at the steady state and inflamed conditions (**Fig. 4**, right). In this analysis, CUP5 was the only virulence gene identified as being required for adaptation to the inflamed oral environment compared to the steady state (nOC/iOC ratio). In the gut, Chu, T6SS1, Fim, CUP5, and CUP6 were identified as required genes for adaptation to the inflamed environment (nGT/iGT ratio). Moreover, several virulence factors were found to be required for optimal ectopic colonization in the gut but not for colonization in the original niche – the oral mucosa. These results showed that Ka SK431 uses more complex virulence mechanisms to adapt to the distant mucosal site – the gut.

### Identification of additional fitness genes required for the colonization in the oral and gut mucosae

By far, we found that several virulence genes associated with iron acquisition, competition, and adhesion are required for oral and gut colonization. Next, we analyzed the requirement of additional genes (i.e., virulence factors other than iron acquisition, competition, and adhesion and non-virulence genes) in in vivo adaptation of SK431. We found various additional genes essential for the in vivo colonization of *K. aerogenes* SK431. **S2 Fig** lists 30 genes required for colonization in the inflamed gut mucosa. To address whether any of these genes are transcriptionally regulated, we analyzed RNA-Seq data (RNA-Seq analysis to be described below). Only 4 genes identified in the Tn-Seq analysis were significantly increased (ApaG and CutA) or decreased (Ribosomal RNA and LdrA) at the transcription level (*p* <0.05) in the inflamed gut compared to the non-inflamed gut (**S2 Fig**, right). Thus, in addition to genes associated with iron acquisition, competition, and adhesion, SK431 uses additional mechanisms for adapting the oral or gut environments.

### Pathways regulated transcriptionally in the oral and gut mucosae

To determine if fitness or virulence genes required for the site-specific adaptation of SK431 identified by the Tn-Seq are also regulated at the mRNA expression level, we compared transcriptomic profiles of SK431 in different environments (oral and gut, non-inflamed and inflamed). (**S3 Fig** and **S1 Table**). To obtain enough numbers of sequence reads from the limited numbers of colonizing bacteria, particularly at periodontal sites, we used SK431-monocolonized mice for this experiment. As shown in **S3 Fig. A**, germ-free (GF) mice were inserted with ligatures soaked with SK431. Inserted ligatures were left in place for 1 day (control, non-inflamed oral cavity [nOC]) and 14 days (inflamed oral cavity [iOC]) (**S3 Fig. A**). Consistent with a previous study [17], increased alveolar bone loss at the ligature-installed site was observed which was associated with severe inflammation in the gingival tissues of the iOC group (**S3 Fig. B-E**). In contrast, no overt signs of inflammation were evident in the nOC group (**S3 Fig. B-E**). Likewise, gut inflammation was induced by oral DSS administration. GF mice were orally inoculated and colonized with SK431 for two weeks, and then colitis was induced by administration of 1.5% DSS for five days (inflamed gut [iGT]) (**S3 Fig. A**). The control group did not receive DSS (non-inflamed gut [nGT]). As shown in **S3 Fig. F-H**, DSS administration, but not control water treatment, resulted in the development of a more severe colitis (i.e., an increased level of fecal lipocalin-2 (Lcn2), a non-specific marker for intestinal inflammation, and displayed histological inflammation in the colon). Bacterial RNA was prepared from the ligature (oral) and fecal (gut) samples and used for the microbial RNA-Seq.

We first analyzed the overall gene expression profiles in all 4 different conditions (nOC, iOC, nGT, and iGT). A non-metric multidimensional scaling (NMDS) analysis of Bray-Curtis dissimilarity indexes showed that the overall gene expression profile was significantly different between groups in the oral and gut sites with and without inflammation (**Fig. 5A**, *p* <0.05 between two groups in PERMANOVA). Next, we investigated the pathways that are differentially regulated at the transcriptional levels in the different environments (**Fig. 5B**). We found that the expression of genes associated with quorum sensing and biofilm formation, nitrogen metabolism, 1, 3-DAP, and AA (lysine/threonine/methionine/cysteine, K/T/M/C) metabolism were significantly higher (SNR > 1, fold difference >2, FDR< 0.05) in the oral mucosa compared with the gut mucosa at the steady state (nOC vs nGT). In contrast, the expression of genes of CU and Type IV pili was significantly higher in the gut mucosa than the oral mucosa in the absence of inflammation. During inflammation, the expression of genes associated with quorum sensing and biofilm formation, and siderophores was significantly higher in the oral mucosa compared to the gut mucosa (iOC vs iGT). In the inflamed gut (iGT), Ka SK431 expressed genes associated with the metabolism of triacylglycerols, Fat acid (FA), lipids, and isoprenoids, and T1SS (SNR < -1, fold difference >2, FDR< 0.05). The mRNA expression levels of Tat, one of Sec-independent protein translocase, were significantly higher in the inflamed gut (iGT) than in the inflamed oral mucosa (iOC). Of note, none of the functional pathways were significantly different between the steady state and during inflammation at each mucosal site (nOC vs iOC and nGT vs iGT). Thus, genes associated with some pathways are differentially regulated transcriptionally in different mucosal sites.

**Fig 5.**
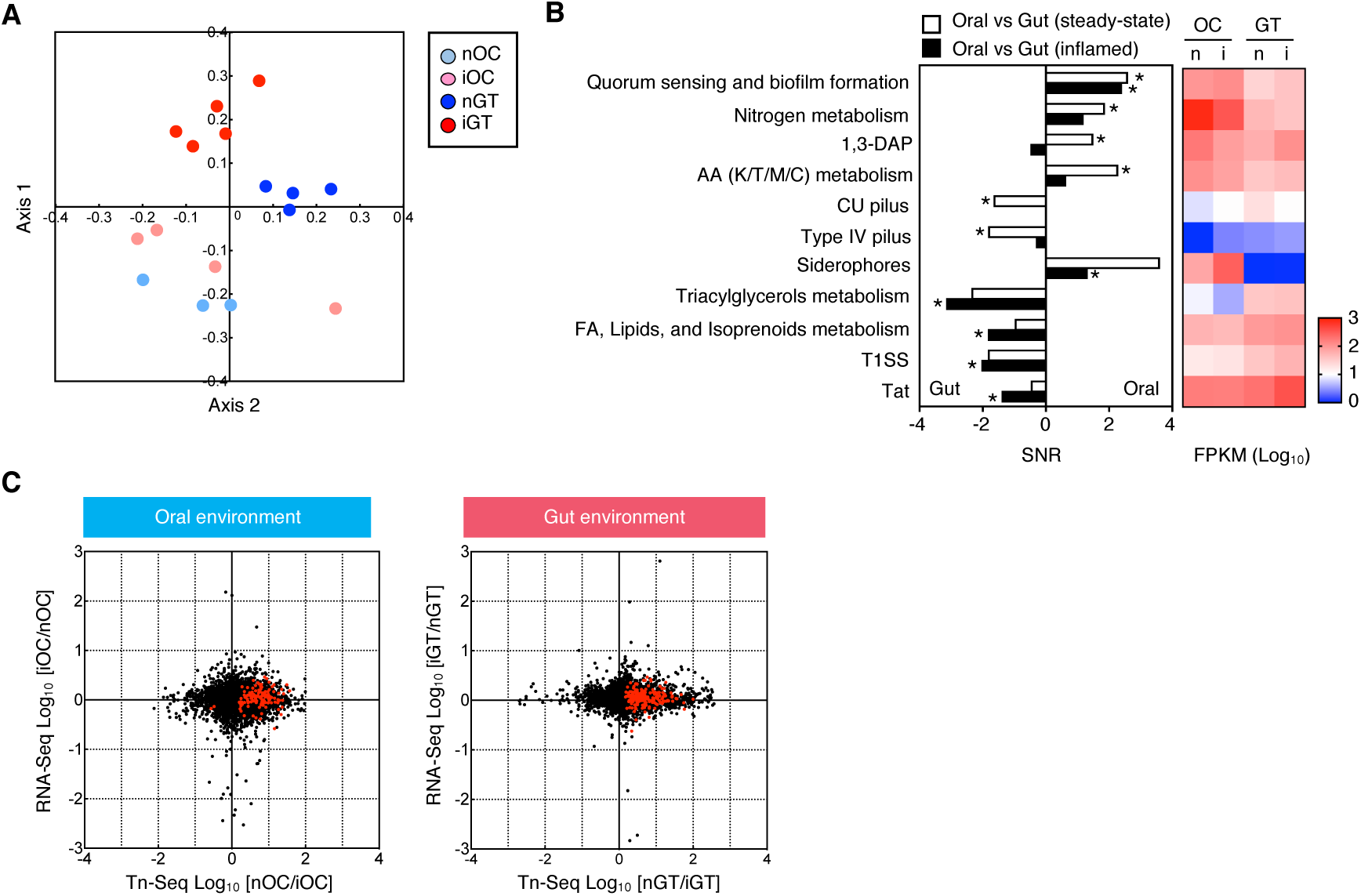
Functional pathways differentially expressed in the oral and gut microenvironments. (A) NMDS plot of Bray-Curtis dissimilarity indexes between groups in the oral and gut microenvironments. (B) Genes were grouped by functional categories and differentially expressed pathways (fold differences > 2, |SNR| > 1, FDR < 0.05, *P* < 0.05) are shown with SNR, which is positive for the ones more expressed in oral cavity. (left). The heatmap on right side shows the different gene abundance (log_10_ averaged FPKM, *P* < 0.05, FDR < 0.05, n = 3 to 4 mice) in the oral and gut mucosae. (C) Gene expression ratio in the oral and gut environments, based on RNA-Seq analyses, compared with gene requirement ratio based on Tn-Seq. Dots indicate the values of individual genes of Ka SK431. Red dots indicate the genes the required genes for the colonization in the oral and gut sites (|SNR| > 1, < 0.05, and FDR < 0.05). *, Pathways with statistically significant difference in mRNA expression levels. CU, chaperone/usher pilus system; FA, fatty acid; FPKM, fragments per kilobase per million mapped fragments; 1, 3-DAP, 1, 3- diaminopropane; T1SS, type I secretion system.

Next, we combined the results of RNA-Seq and Tn-Seq to examine the transcriptional regulation of genes required for oral and gut colonization. In **Fig. 5C**, the gene expression ratio (inflamed/non-inflamed by RNA-Seq) and the gene requirement ratio (non-inflamed/inflamed by Tn-Seq) in each mucosal site (oral or gut) were co-plotted. Although the dominant population of the genes required for colonization in the oral and gut sites were constitutively expressed, particular sets of them were transcriptionally regulated by mucosal site translocation and/or inflammation (red dots, **Fig. 5C**). We further investigated individual genes required for colonization in either the gut or the oral environment and are also transcriptionally regulated at the colonizing sites. To this end, we first selected genes required for the ectopic gut colonization by comparing Tn-Seq results between oral and gut samples (absolute SNR >0.5, fold difference >10, FDR< 0.05 by the Tn-Seq). From identified required genes, we further selected genes expressed at greater levels in the gut than in the oral environment (fold difference >2, FPKM>100 by the RNA-Seq). In this analysis, we identified 19 genes, including EcpA and BglJ, required for the gut, but not oral, colonization that were also transcriptionally regulated (**S4 Fig**). Likewise, we performed the same analysis for oral-specific genes and identified only one gene Fabl, by these selection criteria (**S4 Fig**). These results showed the differential expression of various *K. aerogenes* genes in the oral and gut mucosae at the steady state and during inflammation, further supporting that *K. aerogenes* uses different adaptation mechanisms at these two mucosal sites.

## Discussion

Our previous study reported that periodontal inflammation is a trigger that can induce or exacerbate intestinal inflammation [17]. Oral inflammation results in the expansion of oral pathobionts such as *K. aerogenes* in the oral cavity, which, in turn, are ingested and translocated to the gut where they cause colitis [17]. In the current study, we identified virulence genes exploited by the oral pathobiont *K. aerogenes* to adapt to the ectopic mucosal site (i.e., the GI tract), particularly during inflammation.

In the mucosal surfaces, such as the oral cavity and the GI tract, numerous numbers of symbiotic bacteria exist that prevent the colonization and/or expansion of exogenous or indigenous pathobionts [45]. Pathobionts, therefore, have evolved strategies to overcome the colonization resistance to proliferate in the mucosal sites. To this end, pathobionts acquired specific virulence attributes that increase the ability to adapt to the colonizing niches and overcome competing symbionts. For this purpose, pathobionts employ several virulence factors required for the tissue adhesion, acquisition of nutrients, such as iron, and competition with other bacteria [46, 47].

Fimbriae are known to play an important role in the adhesion of *Klebsiella* species to host epithelial cells [48]. For example, *K. pneumoniae* forms a biofilm using type 1 and type 3 fimbriae to establish colonization in the bladder during the course of catheter-associated urinary tract infections [49, 50]. In this study, we found that *K. aerogenes* strains harbor various types of fimbriae encoded by the operons *fim*, *cfa*, *ecp*, *cup1* – *cup6*. Among these operons, Fim and Cfa are two common types of fimbriae required for the colonization in both the oral and gut mucosae. In addition to Fim and Cfa, *K. aerogenes* SK431 requires five other fimbriae/pili gene clusters (Ecp, CUP1, CUP3, CUP5, and CUP6) to adapt to the inflamed gut. Notably, Fim genes were identified as essential fimbriae in both the oral and gut mucosae by Tn-Seq, although the transcriptional regulation was different at the two mucosal sites. The expression of Fim genes is markedly higher in the oral cavity than in the gut, suggesting that *K. aerogenes* SK431 may preferentially form a Fim-mediated biofilm when colonized in the oral cavity.

Iron is an essential nutrient for the host and most microbes. Compared with a healthy status (i.e., <10–24 M Fe^3+^ in serum), iron levels can be extremely low in the host during infection through a process known as nutritional immunity [51]. To establish infection, iron acquisition ability in an iron-deficient environment is essential for bacterial pathogenicity. *K. aerogenes* SK431 harbors various types of siderophores (e.g., enterobactin, salmochelin, and aerobactin), which are small molecules that chelate iron and are critical for the growth and replication of *Klebsiella* spp. [52–54]. We found that enterobactin, which has a greater affinity for iron than host molecules such as transferrin and lactoferrin [55], is an essential virulence gene for *K. aerogenes* SK431 in the gut environment, especially during inflammation. Importantly, *K. aerogenes* SK431 also has other siderophores, such as salmochelin, which counteract host-mediated iron starvation (i.e., Lcn2 induction by the host) during inflammation [56–58]. In this regard, fecal Lcn2 level was significantly elevated during DSS-induced colitis (**S3 Fig. H**). Thus, salmochelin may be used by *K. aerogenes* SK431 to overcome this host defense mechanism. Our previous report showed that ectopic gut colonization by *K. aerogenes* SK431 causes colitis with intestinal bleeding. Consistent with this finding, in the current study, genes related to the uptake of heme (i.e., the chu operon, which is homologous to the shu locus found in *Shigella* and contains genes required for heme uptake, transport, utilization, and degradation) were found to be essential for this pathobiont to colonize the inflamed gut. These findings suggest that *K. aerogenes* SK431 uses the Chu system in an iron-poor, heme-rich environment [59, 60]. In addition to the chu operon, we found that the Sit system is essential for virulence, enabling *K. aerogenes* SK431 to ectopically colonize the inflamed gut. The Sit system is a common system that exists in most pathogenic enterobacteria, such as *E. coli*, *Salmonella* [61], *Shigella* [62, 63], and *Yersinia* [64], but generally absent in non-pathogenic bacteria. It is encoded by sitABCD and required to transport ferrous iron in low oxygen environments. Of note, these three critical virulence systems (i.e., Sit, Chu, and CUP) are present on a single locus named LIG in *K. aerogenes* strains, including SK431.

In addition to LIG virulence, *K. aerogenes* SK431 uses T6SSs to establish ectopic gut colonization during inflammation. T6SSs were first identified in *V. cholerae* as a syringe-like apparatus anchored within the bacterial cell membrane that serves to inject various effector molecules and toxins into target cells [65, 66]. T6SSs target both other bacteria and eukaryotic cells, suggesting a dual role in bacterial competition and pathogenesis. Based on our bioinformatics analysis, *K. aerogenes* SK431 harbors putative T6SS gene clusters within 2 loci. In *K. pneumoniae*, the T6SS is required for competition within the same bacterial species (i.e., intraspecific competition), between different bacterial species (i.e., interspecific competition), and with fungi (i.e., interkingdom competition) [67]. Further studies are needed to unravel the mechanisms by which the oral pathobiont *K. aerogenes* SK431 exploits its T6SS to compete with gut resident microbes to establish ectopic colonization in the gut. Clearly, *K. aerogenes* SK431 requires multiple virulence mechanisms to adapt to its ectopic microenvironment – the gut mucosa. Notably, not all functional genes required for gut adaptation of *K. aerogenes* SK431 are essential to establish the colonization in its original niche (i.e., the oral mucosa). These results suggest that oral pathobionts use complex virulence mechanisms to adapt to ectopic mucosal sites, such as the GI tract.

Based on the RNA-seq data, it appears that some metabolic and virulence genes in *K. aerogenes* SK431 required for optimal colonization in either the oral and gut mucosae – in both the steady state and during inflammation – are also regulated at the transcriptional level. For example, a type 1 fimbriae pili EcpA, known to be important for binding to host epithelial cells and the early stage of biofilm formation [68], was highly expressed, particularly in the gut, and was also found to be required for the colonization using Tn-Seq analysis in the oral cavity and the gut. Likewise, BglJ, a general transcriptional regulator for virulence genes [69, 70], required for gut colonization of *K. aerogenes* by the Tn-Seq, was more highly expressed in the gut than in the oral site. Moreover, the transcription of some genes was changed during inflammation compared to the steady state.

For example, transcription of ApaG (associated with stress-induced mutagenesis) [71] and CutA (associated with copper tolerance) [72] was significantly increased in the inflamed gut than in the non-inflamed gut. It suggests that *K. aerogenes* SK431 counteracts inflammation-induced stress responses by expressing these molecules in the gut. In contrast, transcription of a ribosomal RNA gene was significantly decreased during gut inflammation, although this gene is required for the colonization in the inflamed gut. Similarly, LdrA was significantly decreased in the inflamed gut compared to the non-inflamed gut. LdrA is known to exert toxicity by inhibiting ATP synthesis in bacteria [73]. Therefore, the reduction of LdrA may, in turn, promote DNA replication, transcription and translation, and eventually cell growth of bacteria in the inflamed gut. Thus, transcriptional regulation of several genes may be required for the ectopic gut colonization by *K. aerogenes*.

It is possible that there are additional functional pathways required for the site-specific adaptation of *K. aerogenes* SK431 that were not identified by Tn-Seq analysis. For example, L-serine catabolism plays a crucial role in the fitness of pathogenic Enterobacteriaceae, such as adherent– invasive *E. coli* (AIEC) and *Citrobacter rodentium*, in the inflamed gut [40]. Of note, L-serine catabolism is regulated by the gene operons Tdc and Sda, which are functionally redundant [40]. One operon can offset the defect in the other operon; therefore, the requirement of L-serine catabolism would not be identified by Tn-Seq analysis. In this regard, this study found that the L- serine catabolism pathway is conserved in the oral pathobiont *K. aerogenes* SK431. The expression of genes related to L-serine catabolism, such as Tdc operon, is markedly higher in the gut compared to the oral environment, and further increased by inflammation (**S1 Table**). This data suggests the possibility that *K. aerogenes* SK431 may use L-serine catabolism in its adaptation to the gut environment, especially during inflammation, as occurs with AIEC [40].

We have demonstrated that the oral pathobiont *K. aerogenes* SK431 exploits distinct virulence and metabolic strategies to establish colonization depending on the environment in which it resides – the oral mucosa or the gut mucosa. This study, however, has limitations. Although we identified virulence and metabolic pathways involved in site-specific adaptations, further functional validations using single mutant strains of the pathobiont are necessary. Also, the possible role of other microbes in the microbiota should be considered, particularly in the transcriptome analysis. For Tn-seq analysis, we used SPF mice. Hence, the fitness genes identified are considered essential in the normal setting (i.e., in the presence of the symbiotic bacteria). In contrast, to obtain sufficient amounts of RNA, we used a *K. aerogenes* monocolonization for RNA-Seq analysis. However, metabolic changes may be influenced by the presence of other microbes. Therefore, the importance of the identified metabolic pathways in the fitness of *K. aerogenes* should be verified in gnotobiotic mice colonized by multiple bacterial species or in conventionally raised animals. Another limitation is related to the bacterial strain used. In this study, we used *K. aerogenes* as a model oral pathobiont, as we and others reported *Klebsiella* spp. reside in the oral cavity in humans and mice, particularly during periodontal disease [16, 17, 21, 22]. However, Enterobacteriaceae such as *Klebsiella* spp. are not considered as classical oral pathobionts of humans and may be enriched only in a subset of patients with oral diseases. In this context, it would be interesting to examine if other oral pathobionts, such as *P. gingivalis*, *F. nucleatum*, and *Prevotella spp.*, which are reported to exacerbate colitis by gavaging orally [13–15], use similar adaptation mechanisms in their ectopic niche. Despite these limitations, this study provides valuable insights, as a proof of concept, into the possible site-specific adaptation mechanisms used by oral pathobionts.

## Author contributions

Y.G. and N.K. conceived and designed the experiments with help from S.K., G.C.-F., D.W., K.S., G.N., and C.J.A.. N.I. performed Tn-seq and RNA-seq analyses. Y.G., N.I., and N.K. wrote the manuscript with contributions from all authors.

## Acknowledgments

The authors wish to thank the University of Michigan Center for Gastrointestinal Research (NIH 5P30DK034933), and the Host Microbiome Initiative, the Germ-Free Mouse Facility, the Advanced Genomics Core for technical assistance, and the In-Vivo Animal for histological evaluation, all at the University of Michigan.

## Data availability

The *Klebsiella aerogenes* SK431 genomic sequence, Tn-Seq and RNA-Seq data used in this study are available from the BioProject website (https://www.ncbi.nlm.nih.gov/bioproject/), accession numbers PRJNA763502.

## Supporting information

**S1 Fig.**
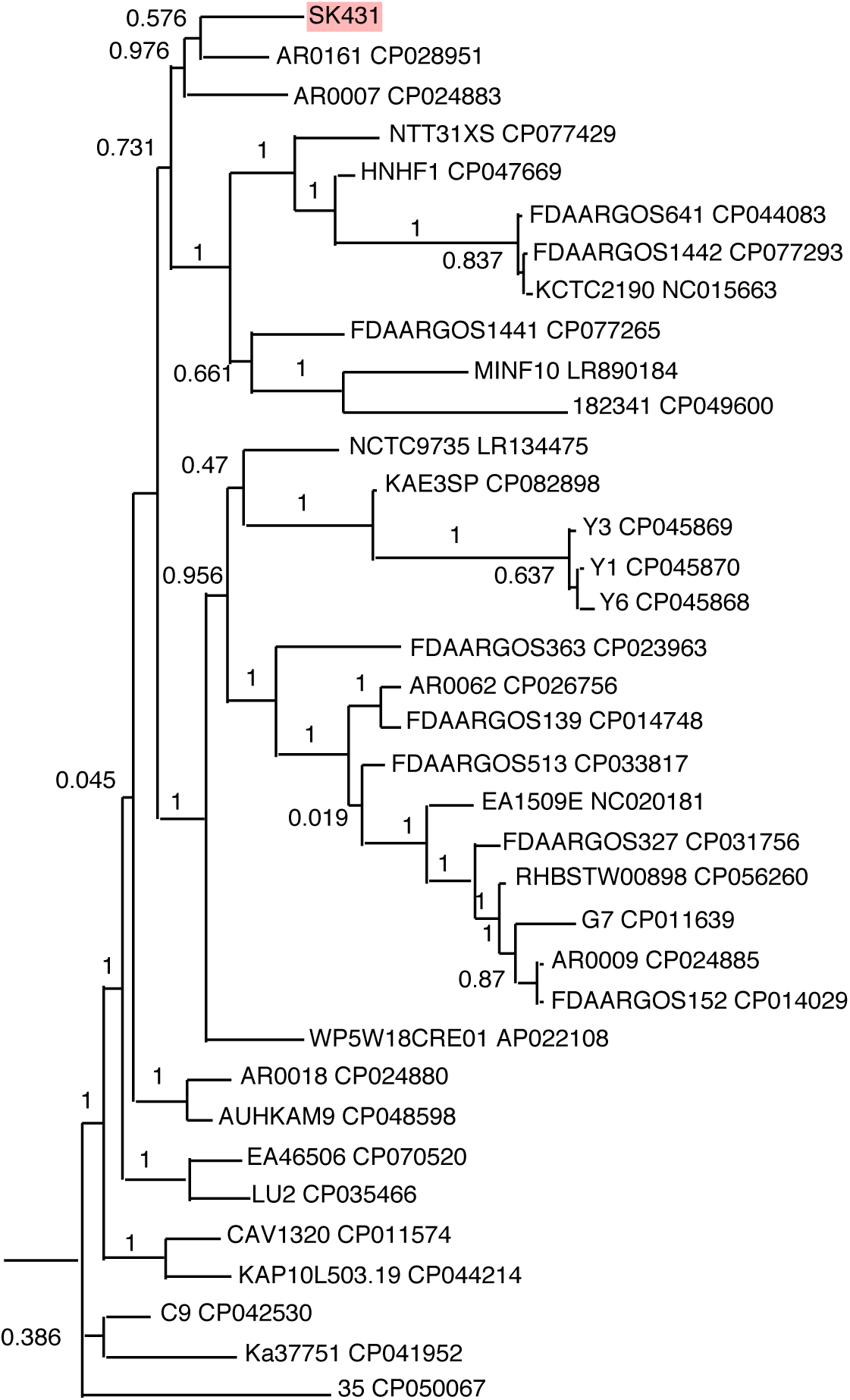
Phylogenetic tree of *Klebsiella aerogenes* strains. The phylogenetic tree of *K. aerogenes* SK431 (highlighted) and 35 reference *K. aerogenes* strains (Strain name + GenBank accession numbers). *K. aerogenes* strains that have complete genomic sequence were selected from this panel and shown in Fig. 1

**S2 Fig.**
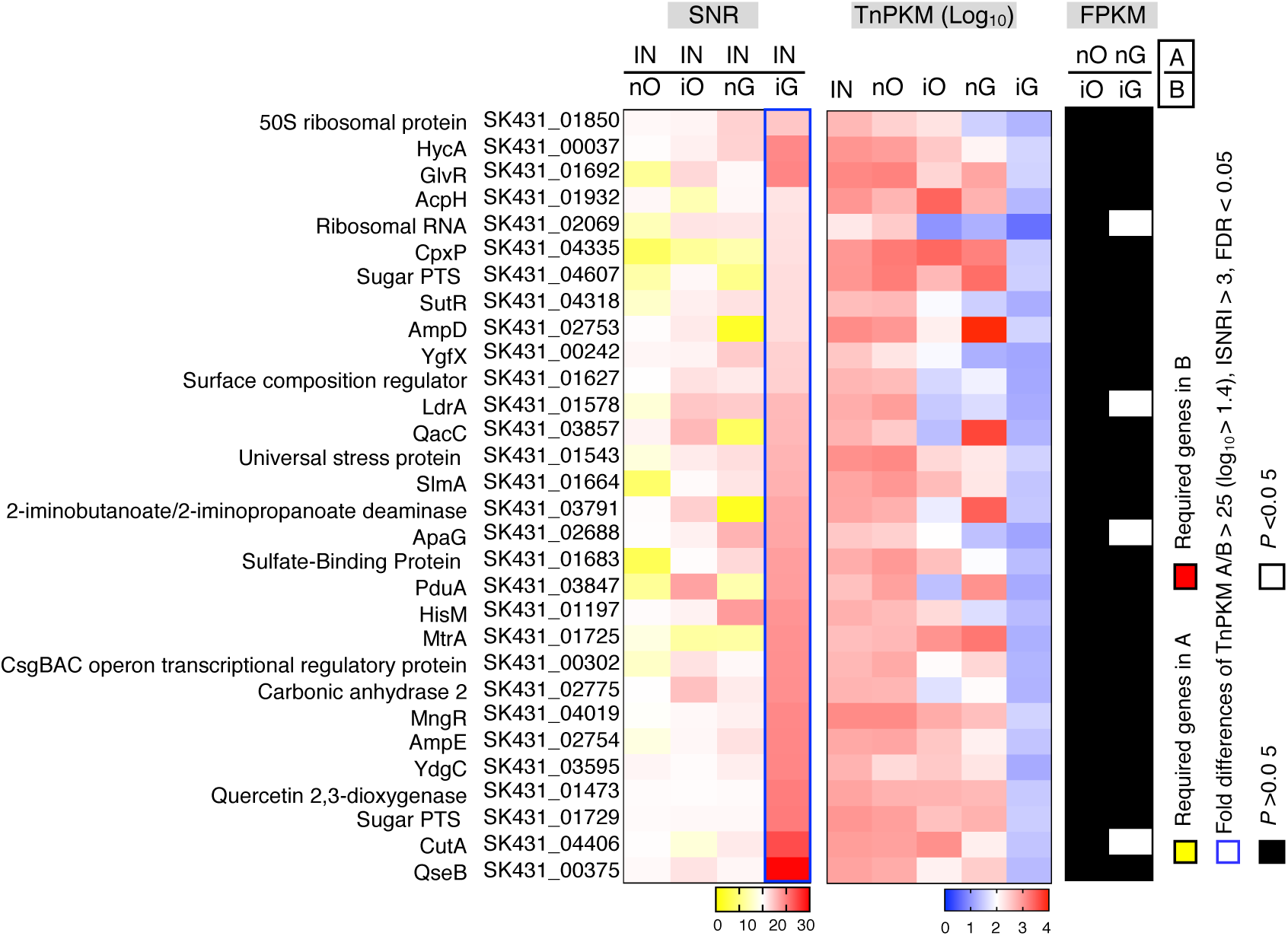
Additional genes required for *K. aerogenes* SK431 colonization at distinct mucosal sites. The relative abundance of gene mutants between input and outputs was analyzed by SNR (left, fold difference >25, |SNR| >3, *P* <0.05, FDR< 0.05). The heatmap (middle) shows the transposon mutant gene abundance (TnPKM) in input (IN) and outputs (nOC, iOC, nGT and iGT). The significantly difference on mRNA expression of non-virulence genes in non-inflamed and inflamed sites (right, fold difference >2, *P* < 0.05).

**S3 Fig.**
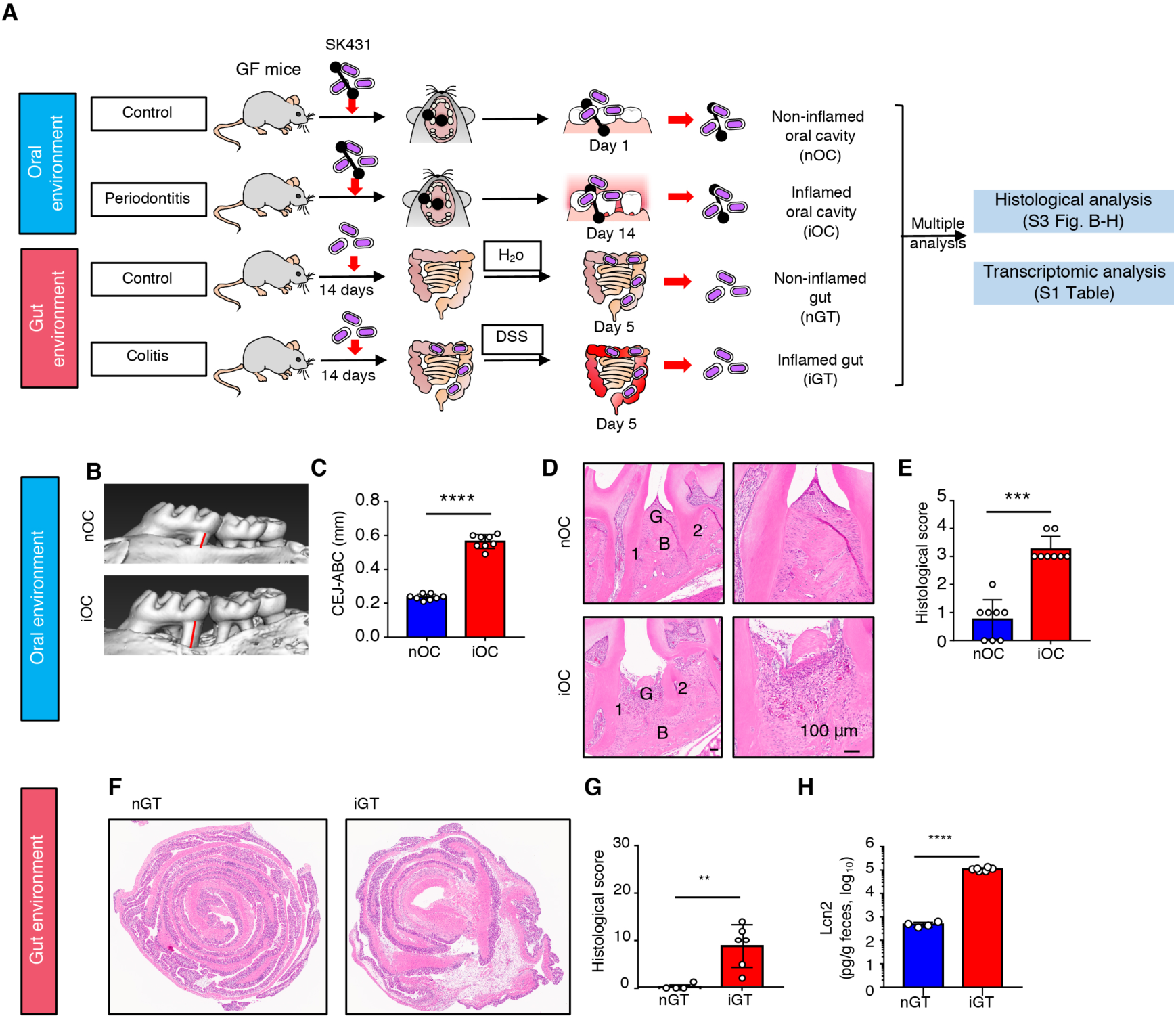
Experiment procedure for RNA-Seq analysis of *K. aerogenes* moncolonized mice from oral and gut mucosae. (A) Experimental design for transcriptomic analysis of the oral pathobiont *K. aerogenes* (Ka) at different digestive organ sites: i) mouth: Germ-free (GF) C57BL/6 mice were inoculated with Ka mutant library-soaked ligature. Oral ligatures were harvested 1 day (control) and 14 days (inflamed) post insertion. The maxilla samples were used to evaluate oral inflammation, ligatures were harvested for bacterial RNA extraction. ii) gut: GF C57BL/6 mice were inoculated with Ka with ligatures (ligatures were pre-soaked in 10^9^ CFU/ml Ka suspension) and colonized for 14 days. Colitis was induced by the administration of dextran sulfate sodium (DSS) for 5 days. Mice were sacrificed, and the colonic tissues were collected to evaluate gut inflammation. Feces were used for bacterial RNA isolation. (B–E) Mice moncolonized with SK431 developed severe periodontitis after ligature. (B) Representative micro-CT images of the alveolar bone. (C) The distance (indicated by bars) from the cementoenamel junction (CEJ) to the alveolar bone crest (ABC) on the buccal side of the ligature site. (D) Representative HE-stained sections of the gingival tissue. 1, root of the first molar; 2, root of the second molar; B, alveolar bone; G, gingival epithelium. (E) Histological score of the gingival tissue (to a maximum of 4). (F–H) Mice moncolonized with SK431 developed colitis after DSS treatment. (F) Representative histological images of the colonic tissue. (G) Histological score of the colonic tissue (to a maximum of 32). (H) Fecal lipocalin-2 (Lcn2) level. (B, D, E, G) Results are the mean ± SD. (oral: n = 8 [control, periodontitis], gut: n = 4 [control]; n = 6 [DSS]). \*\**P* < 0.01, \*\*\**P* < 0.001, \*\*\*\**P* < 0.0001 by Student *t* test (for C and H) or Mann-Whitney *U* test (for E and G).

**S4 Fig.**
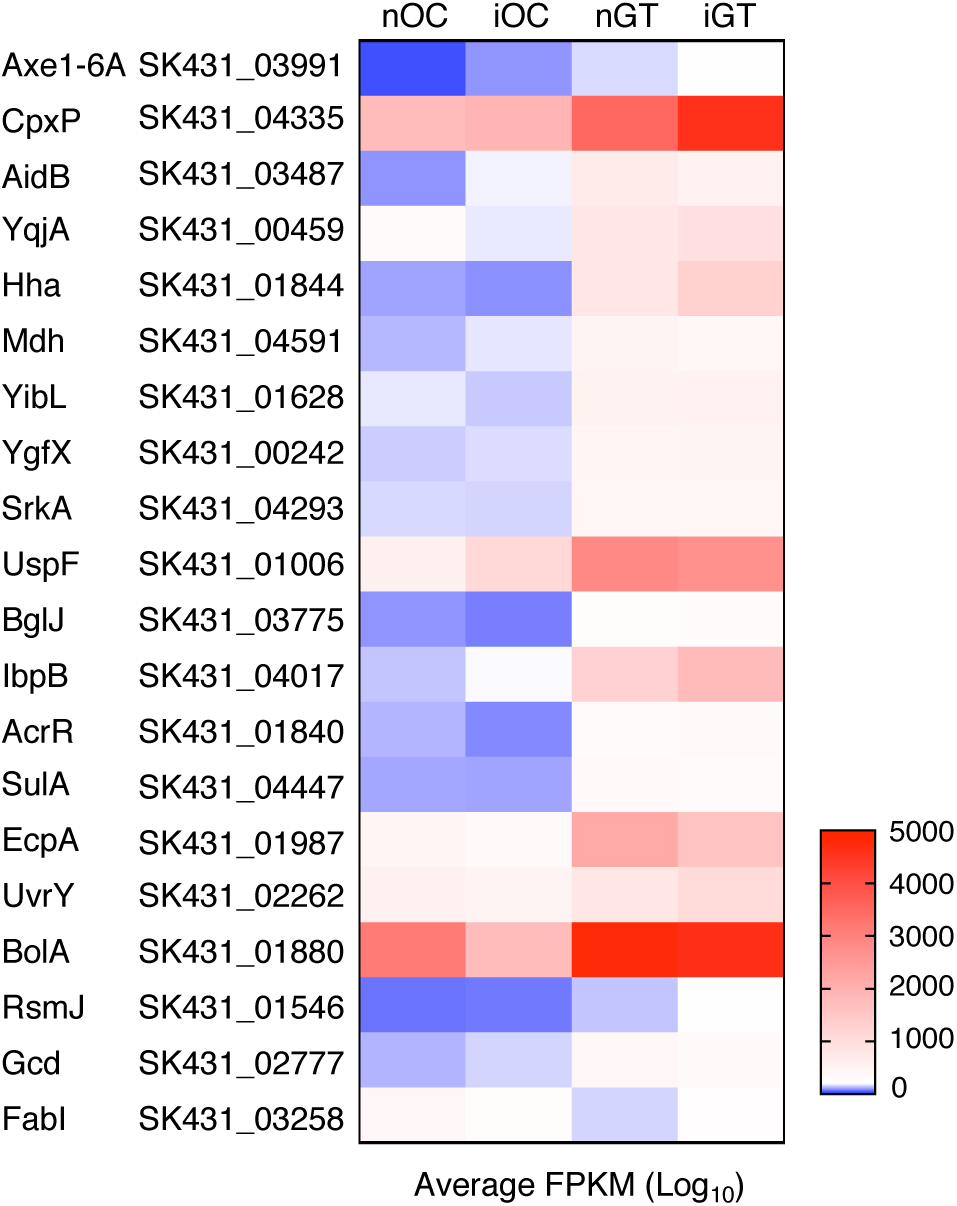
Site-specific expression of virulence and metabolic genes in *K. aerogenes* SK431. mRNA expression levels (log_10_ averaged FPKMs) at the two different mucosal sites – oral and gut – in the steady state and during inflammation. The genes listed here were identified as the required gene for the colonization in either the gut or the oral environment (fold difference >10, |SNR| >0.5, FDR< 0.05 by the Tn-Seq) and are also transcriptionally regulated at the colonizing sites (fold difference >2, FPKM>100 by the RNA-Seq).

**S1 Table.**
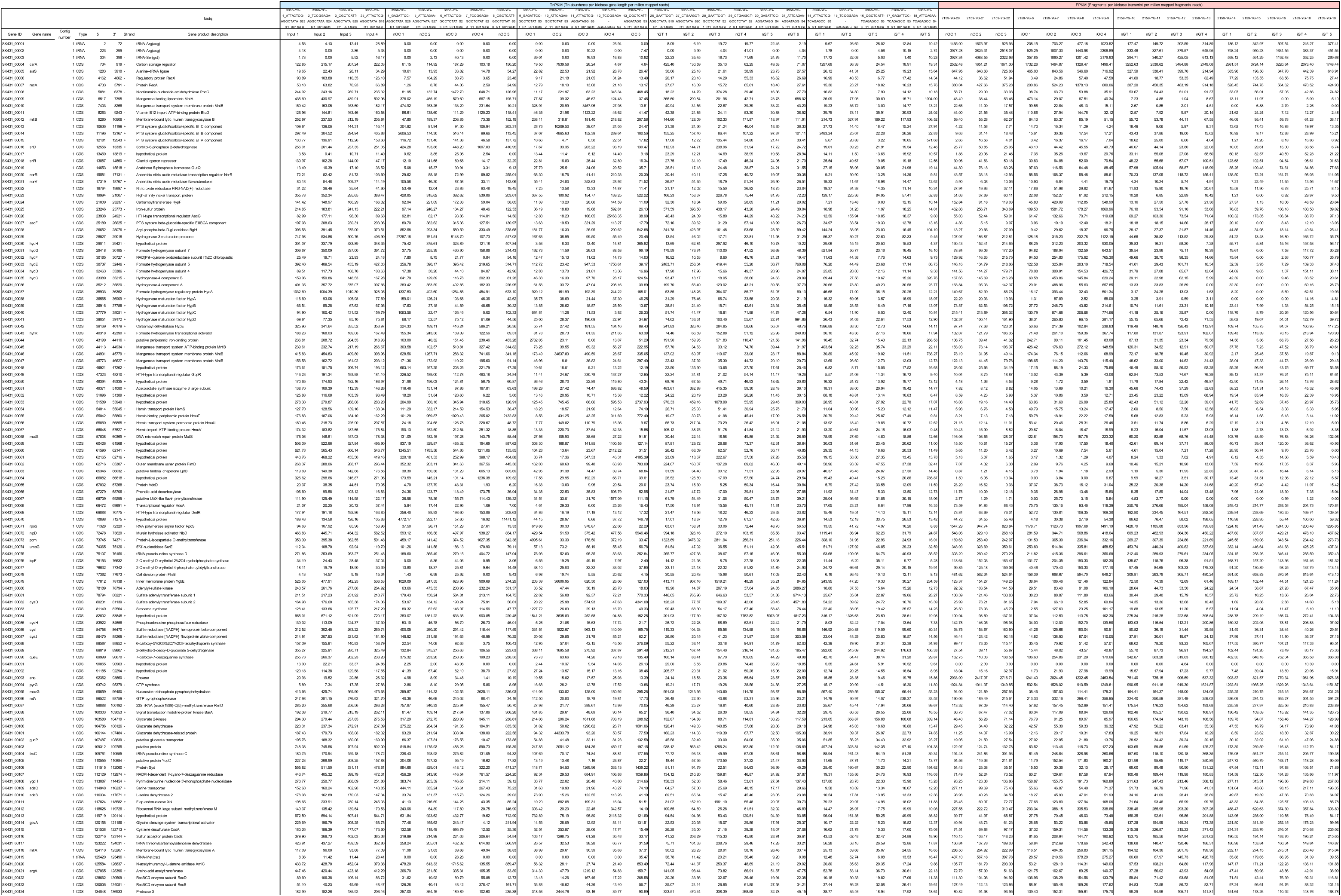
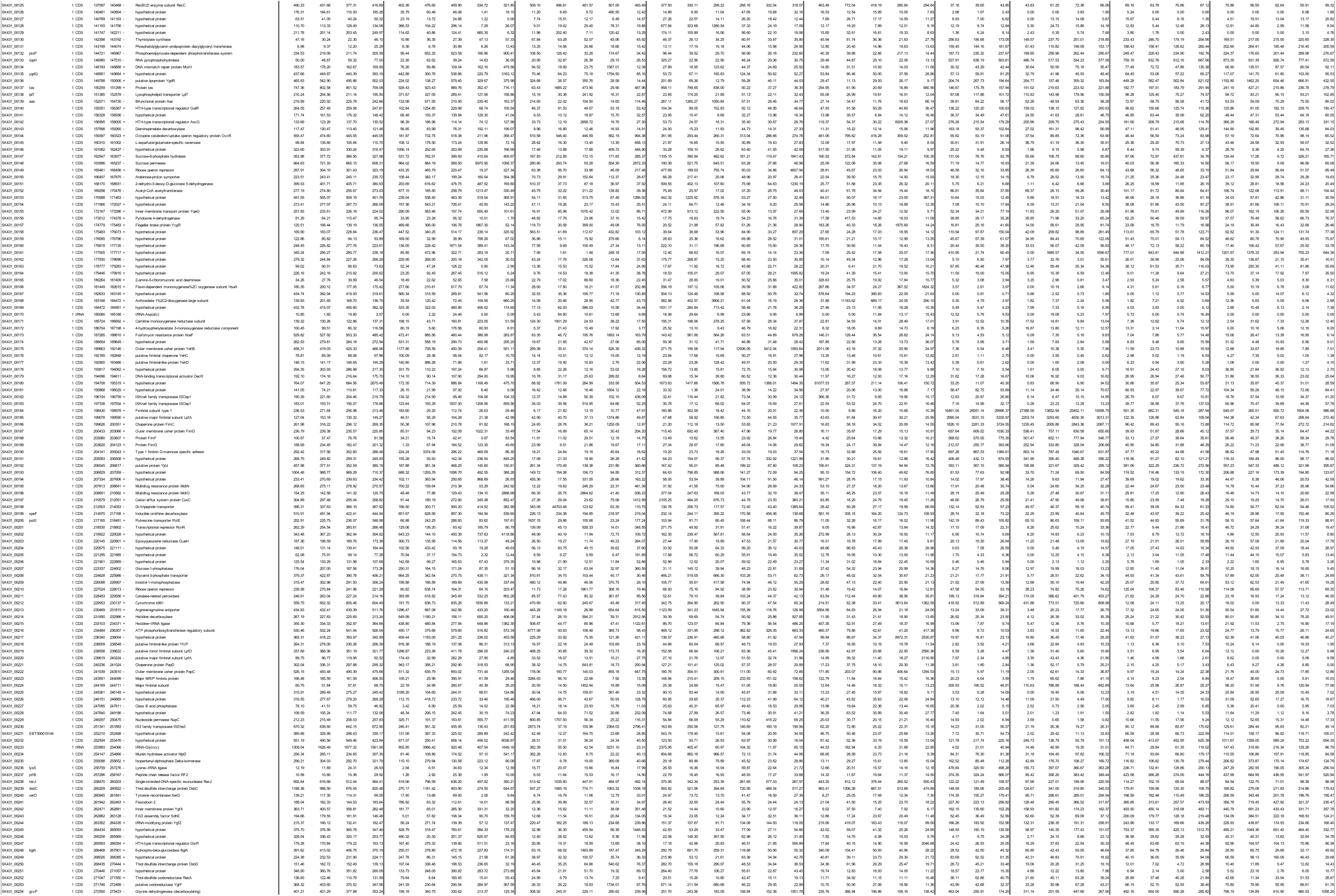
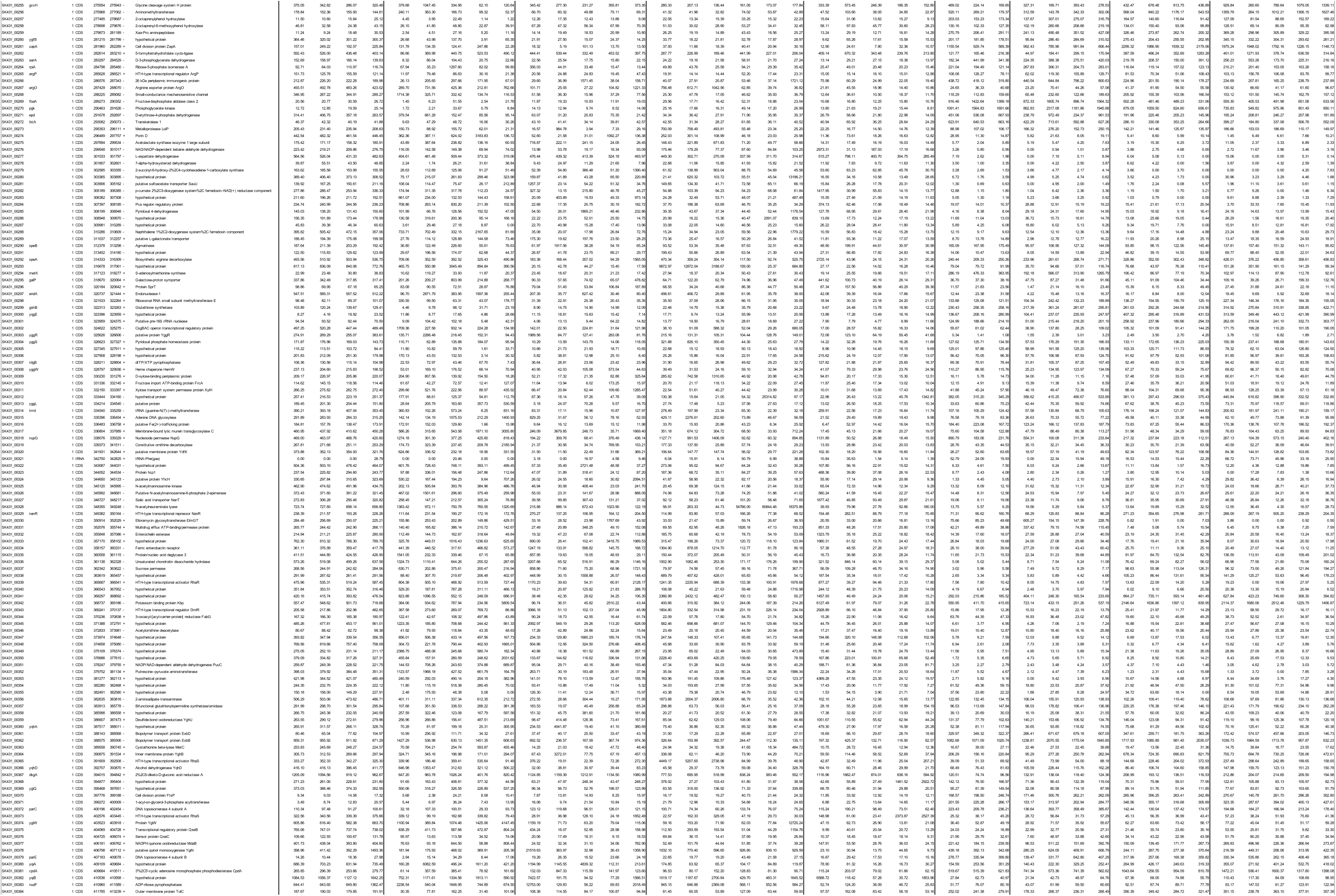
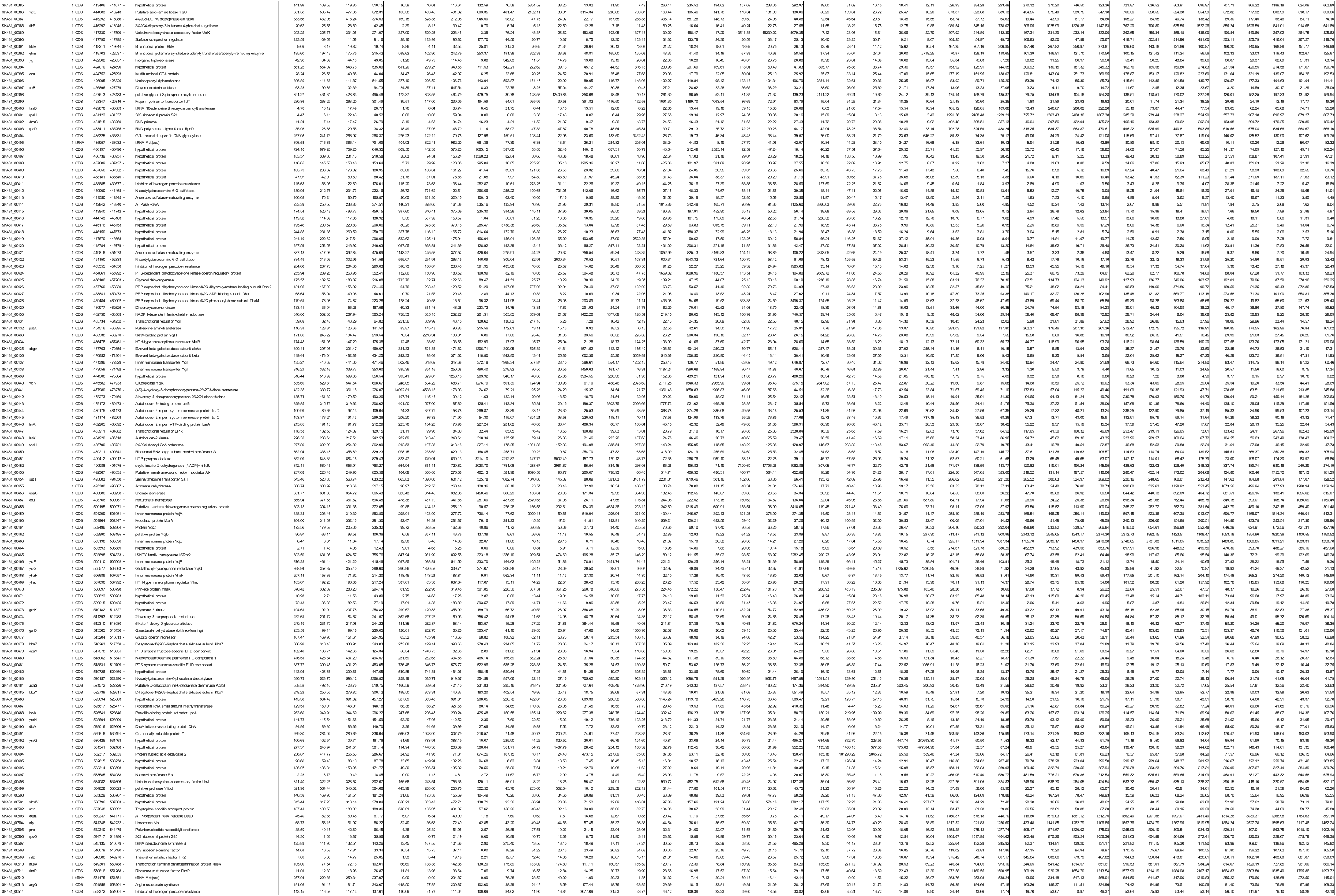
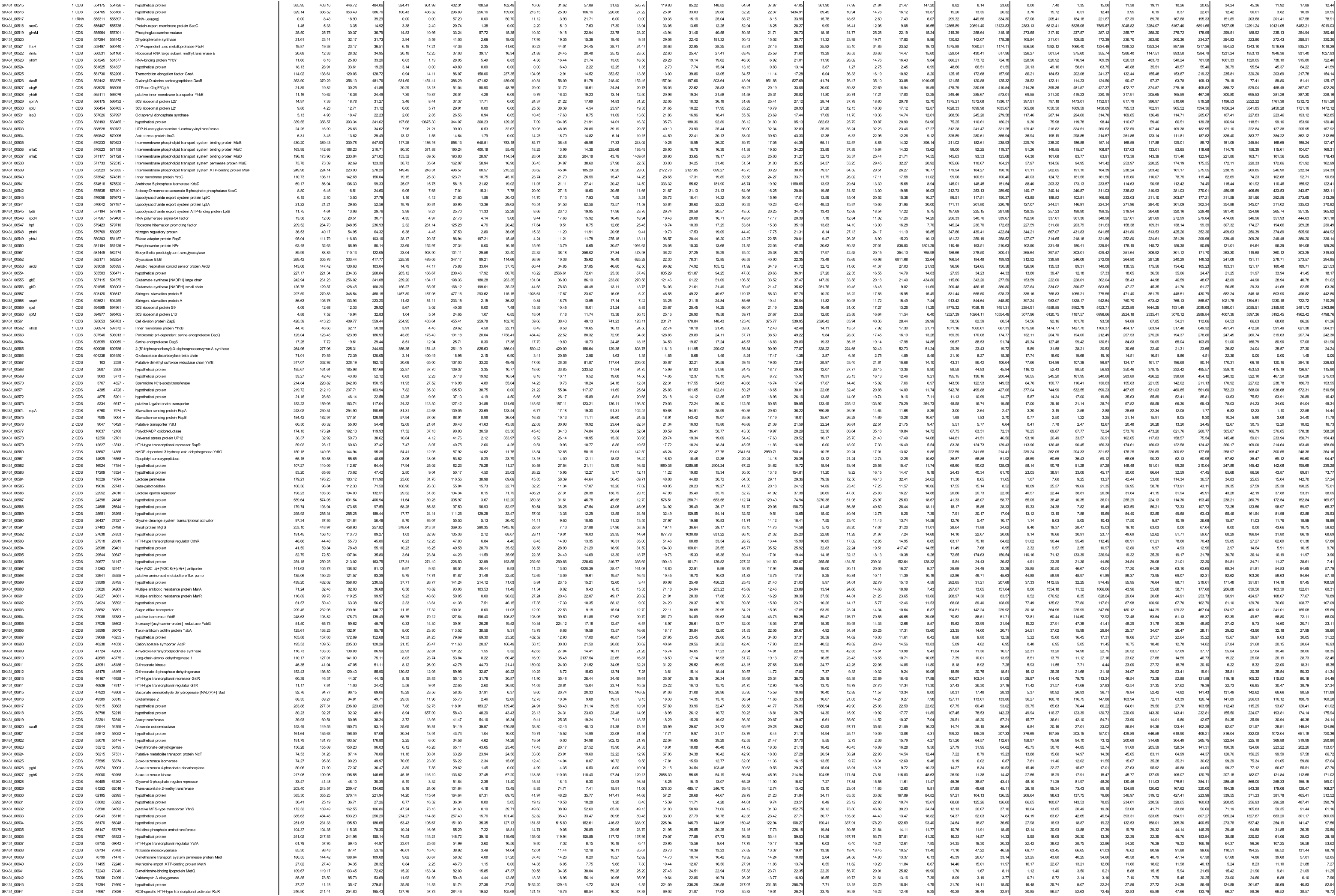
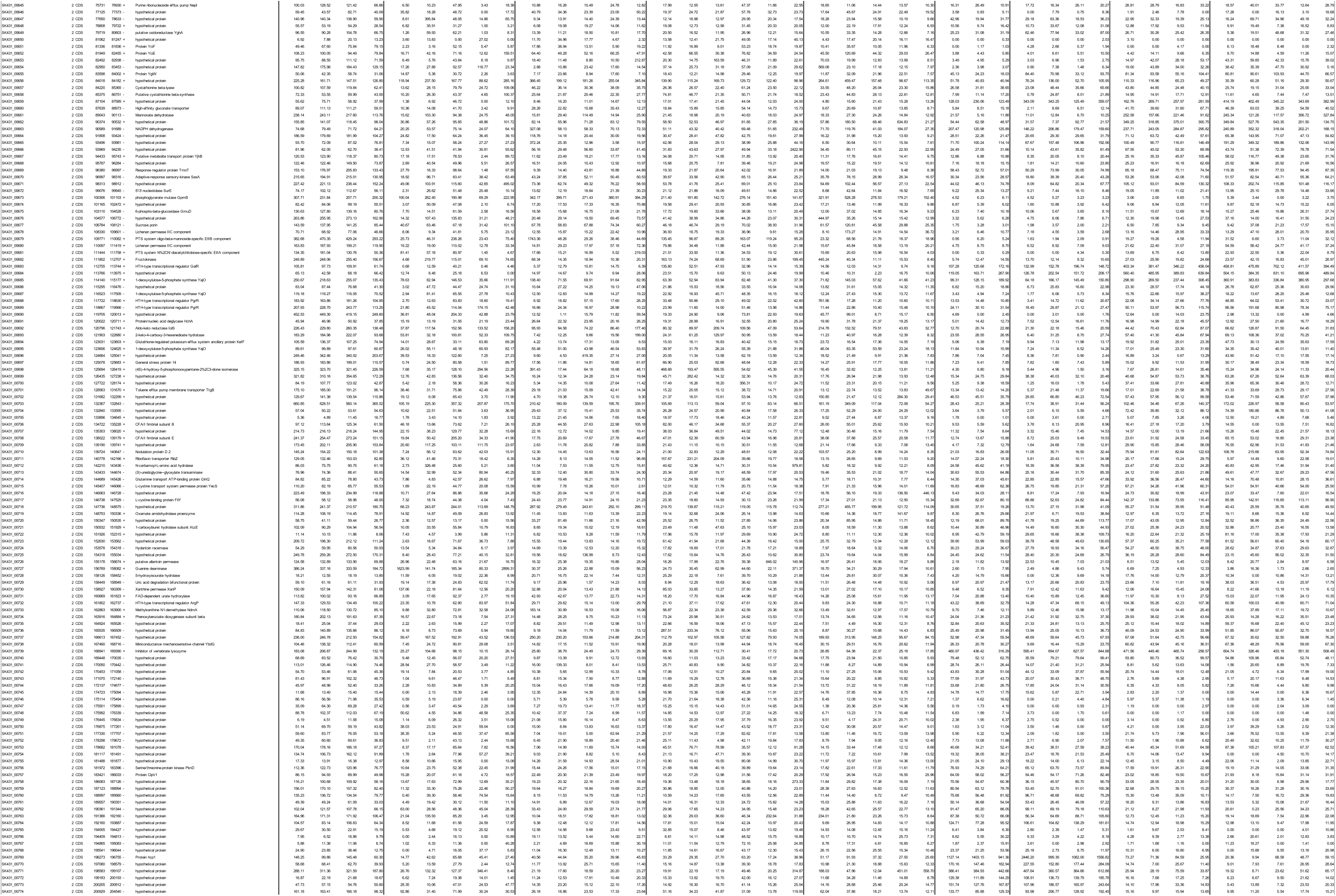
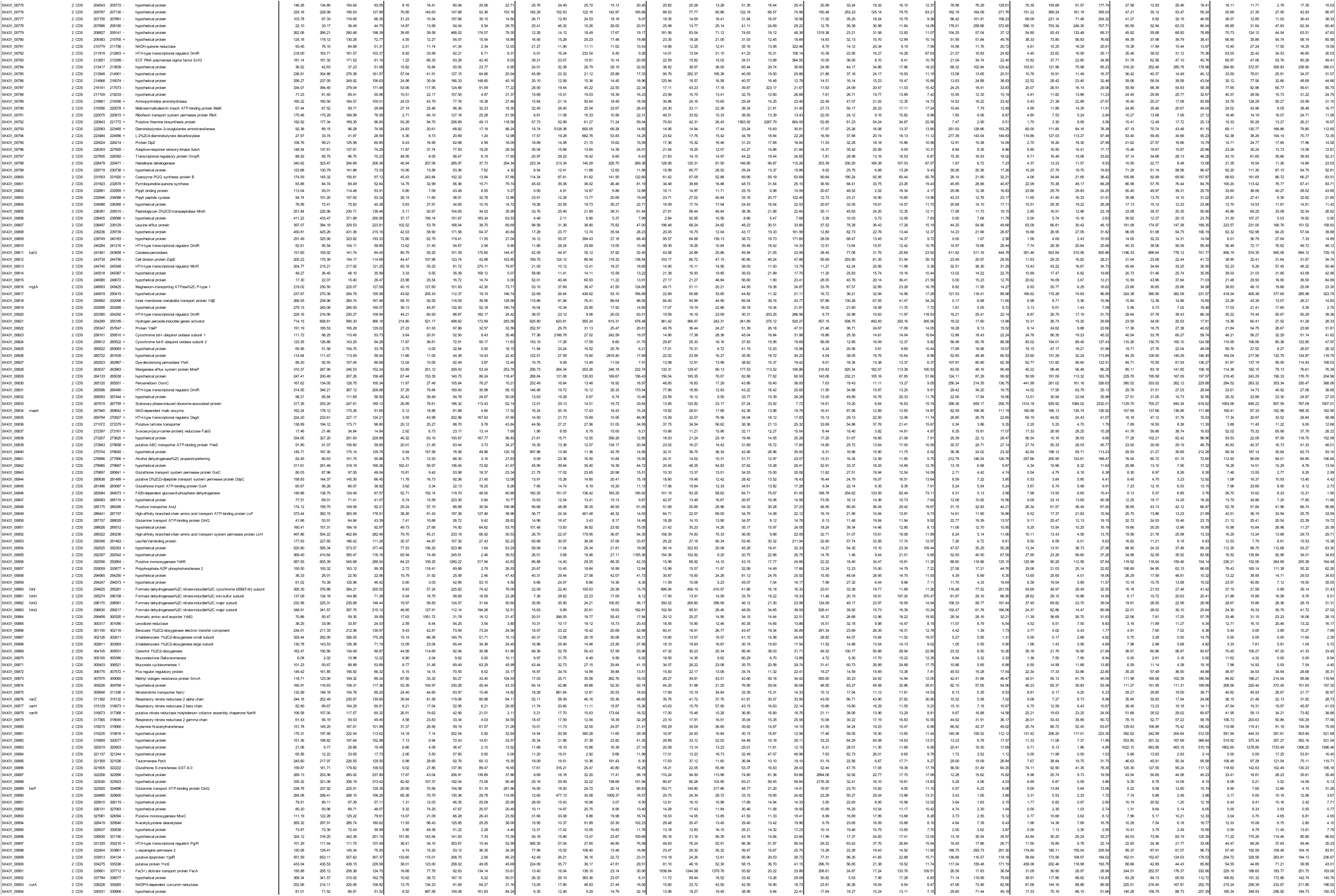
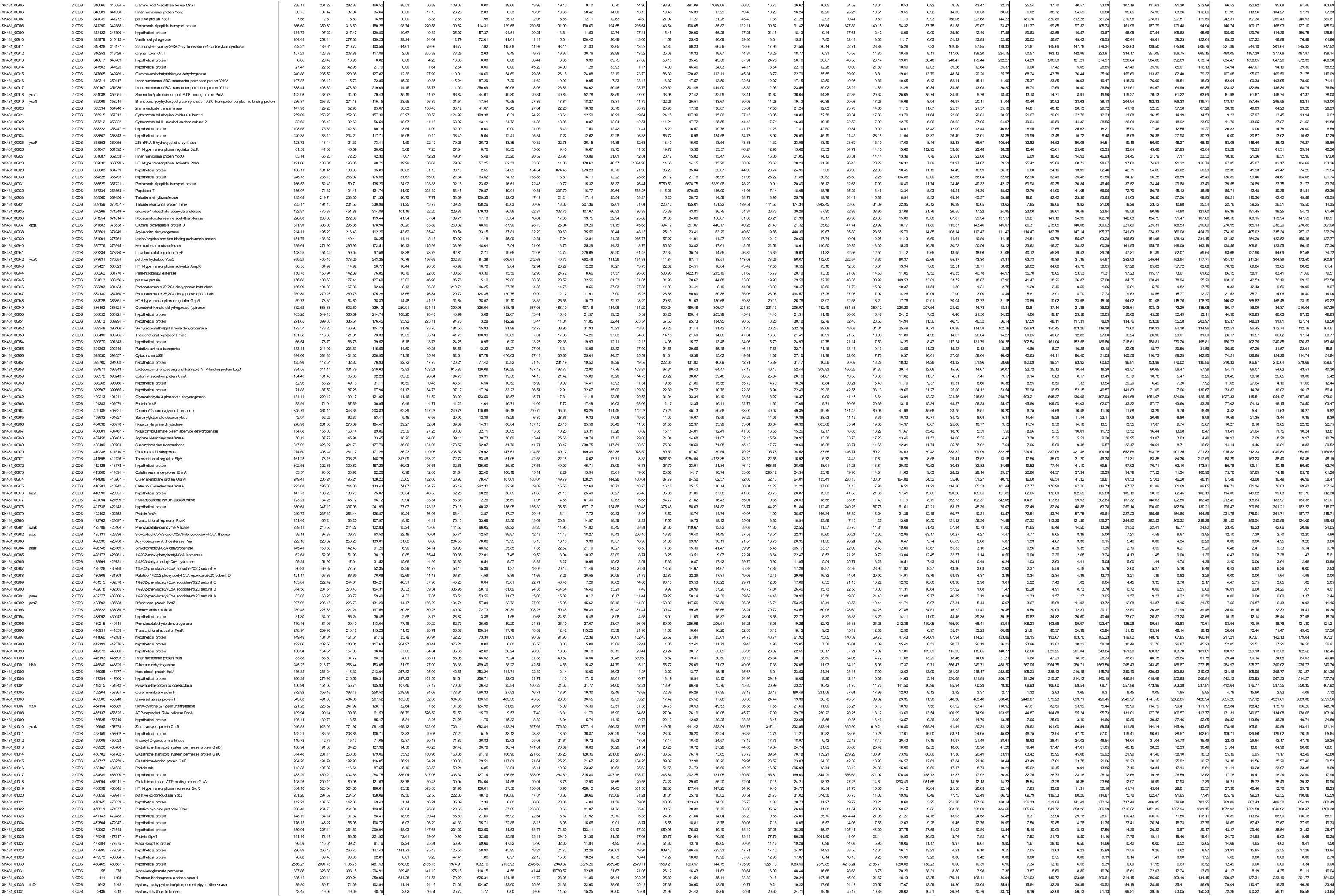
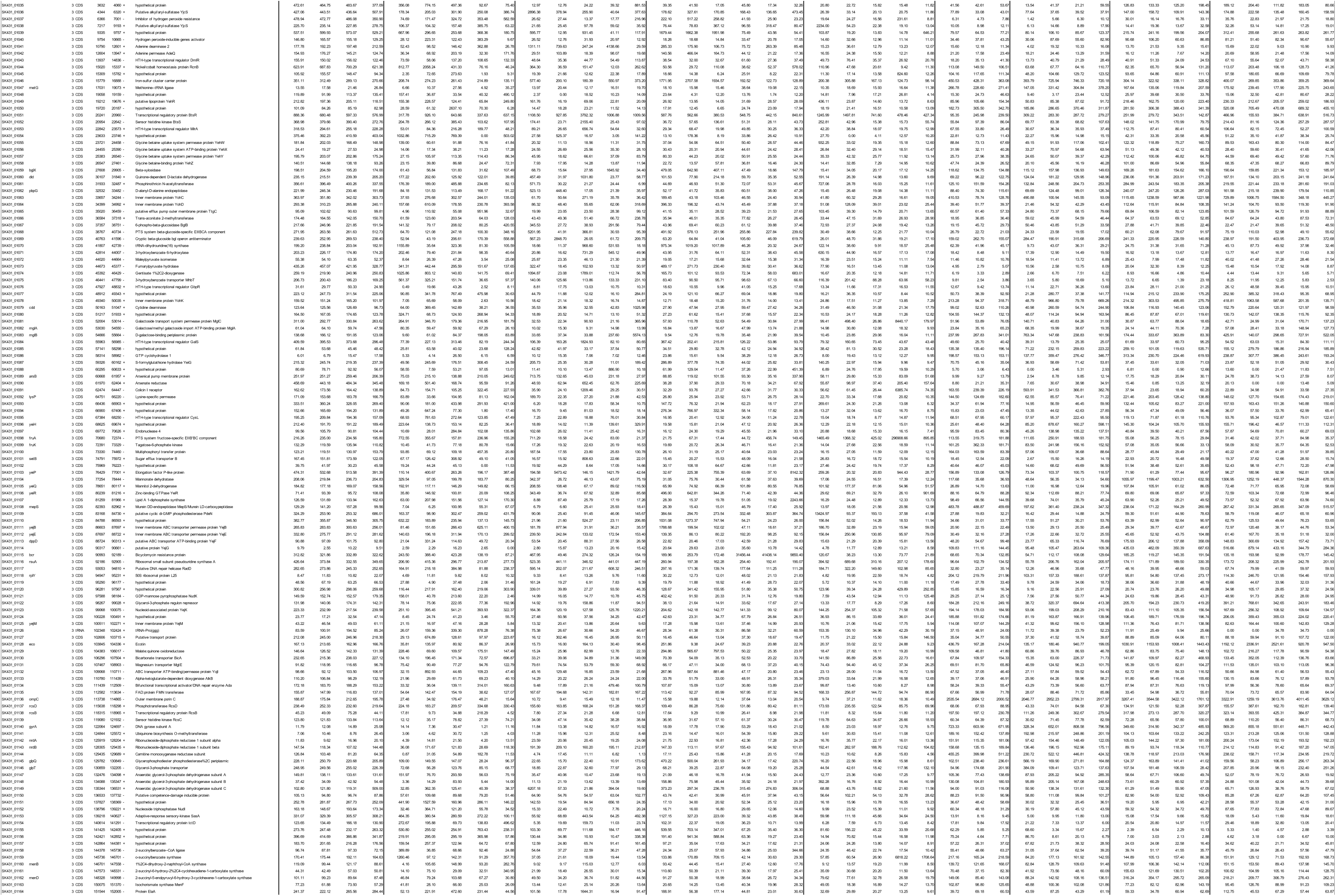
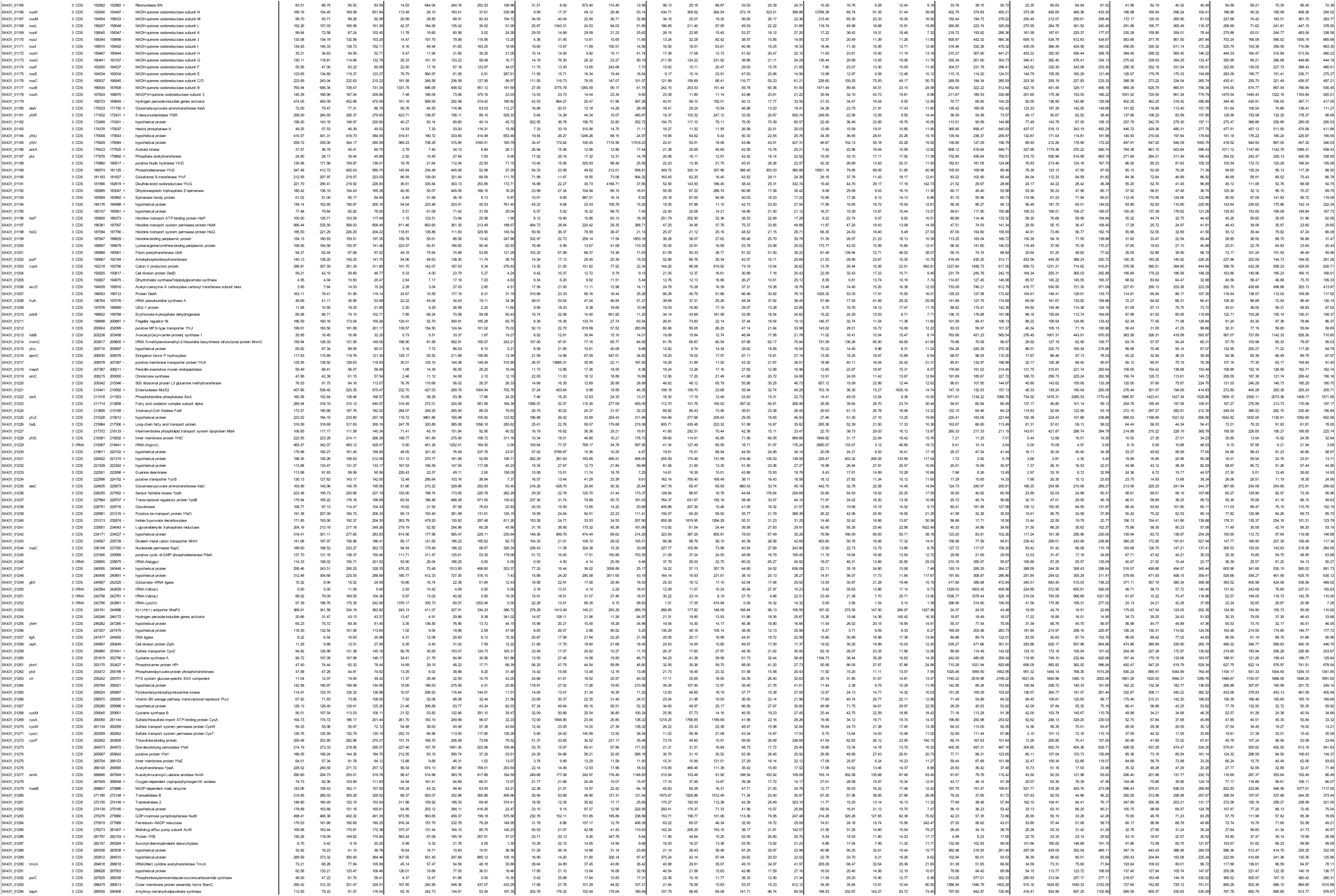
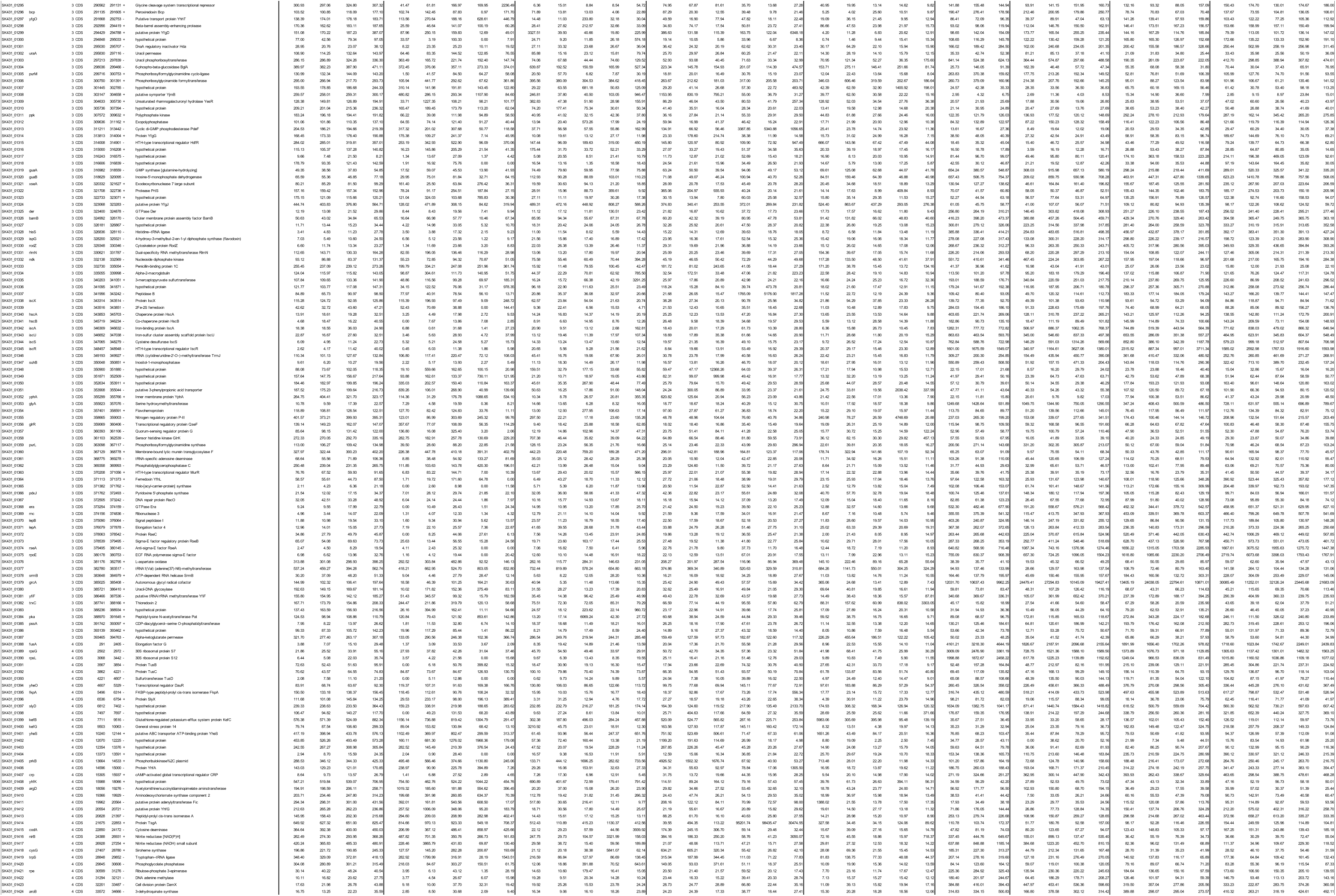
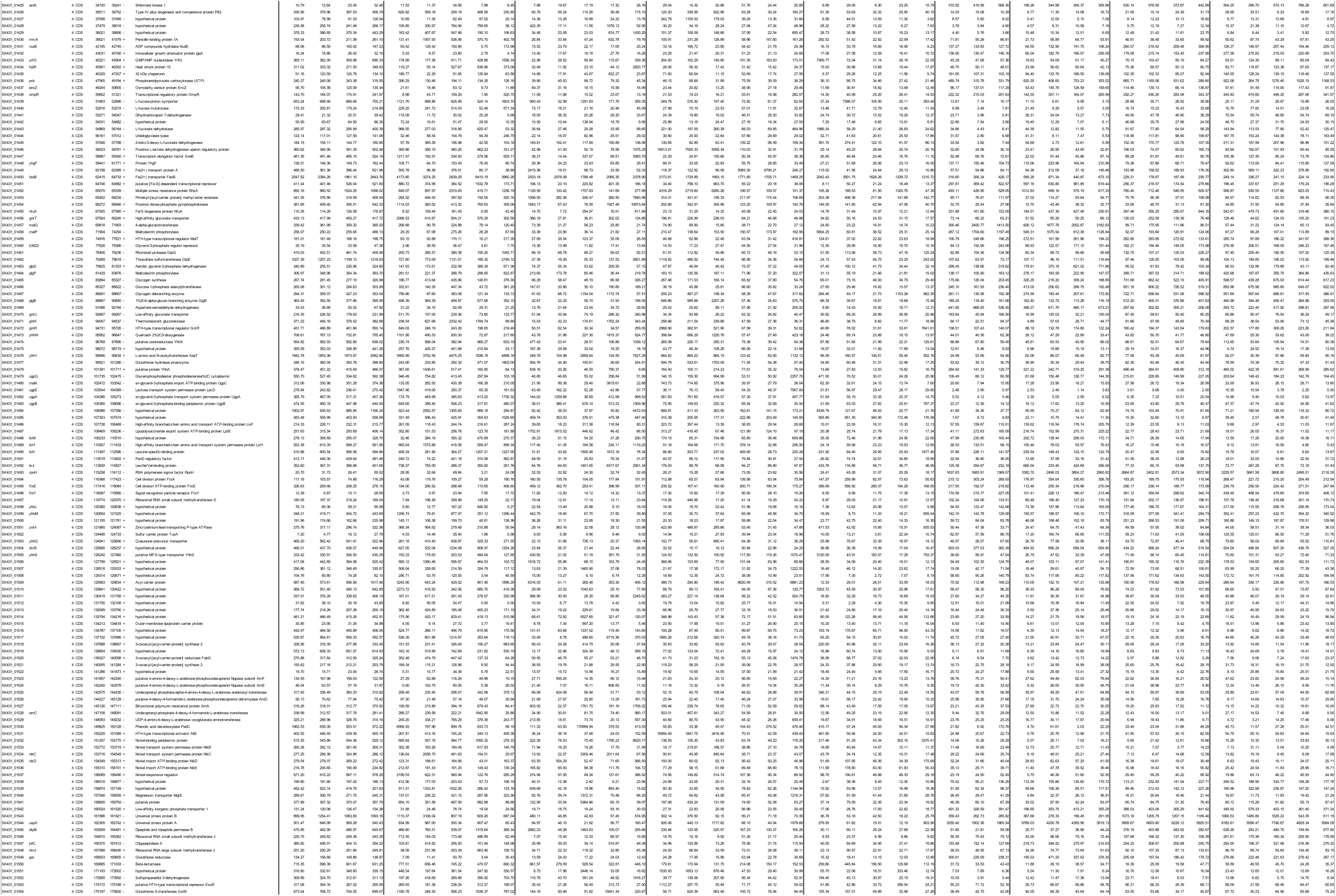
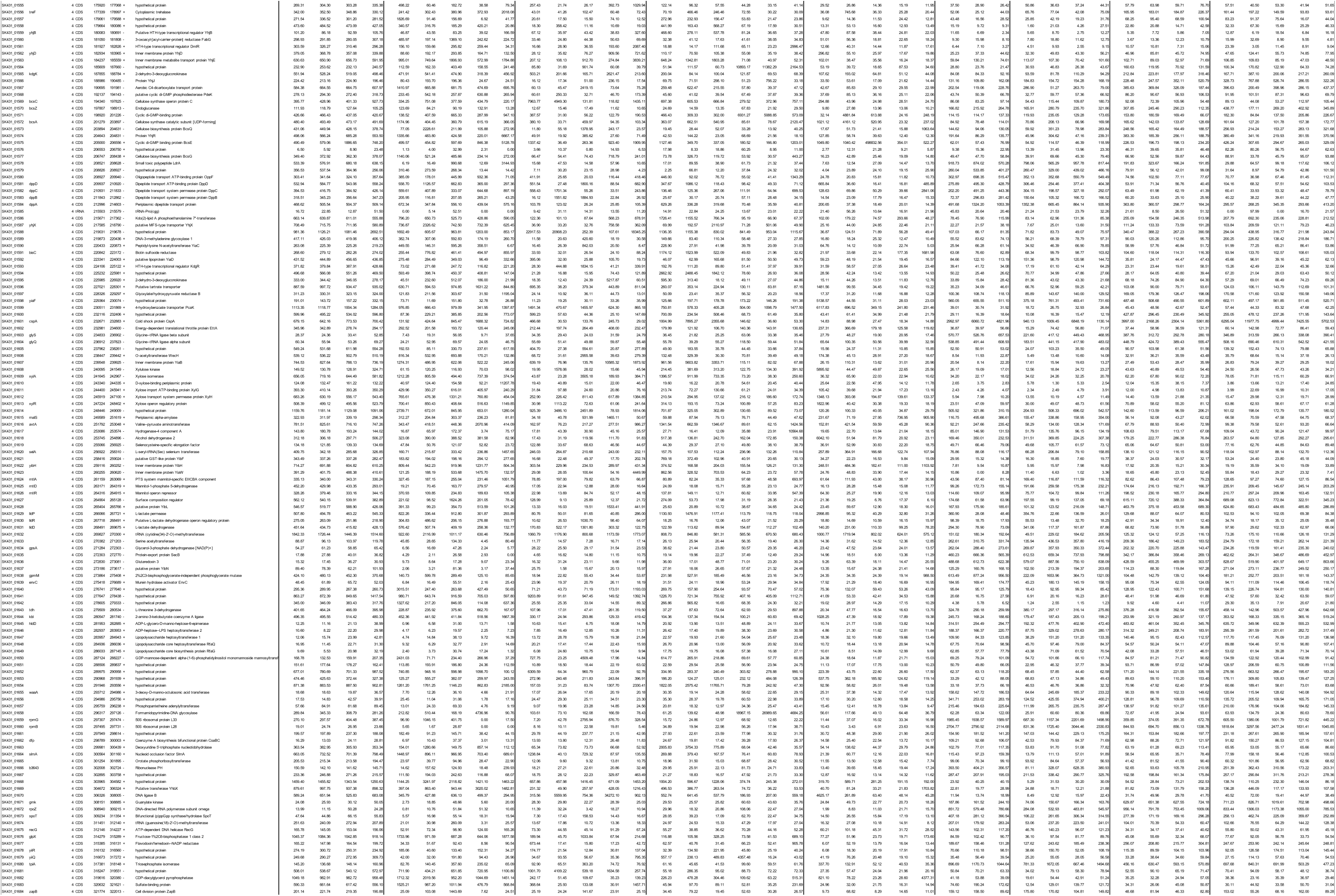
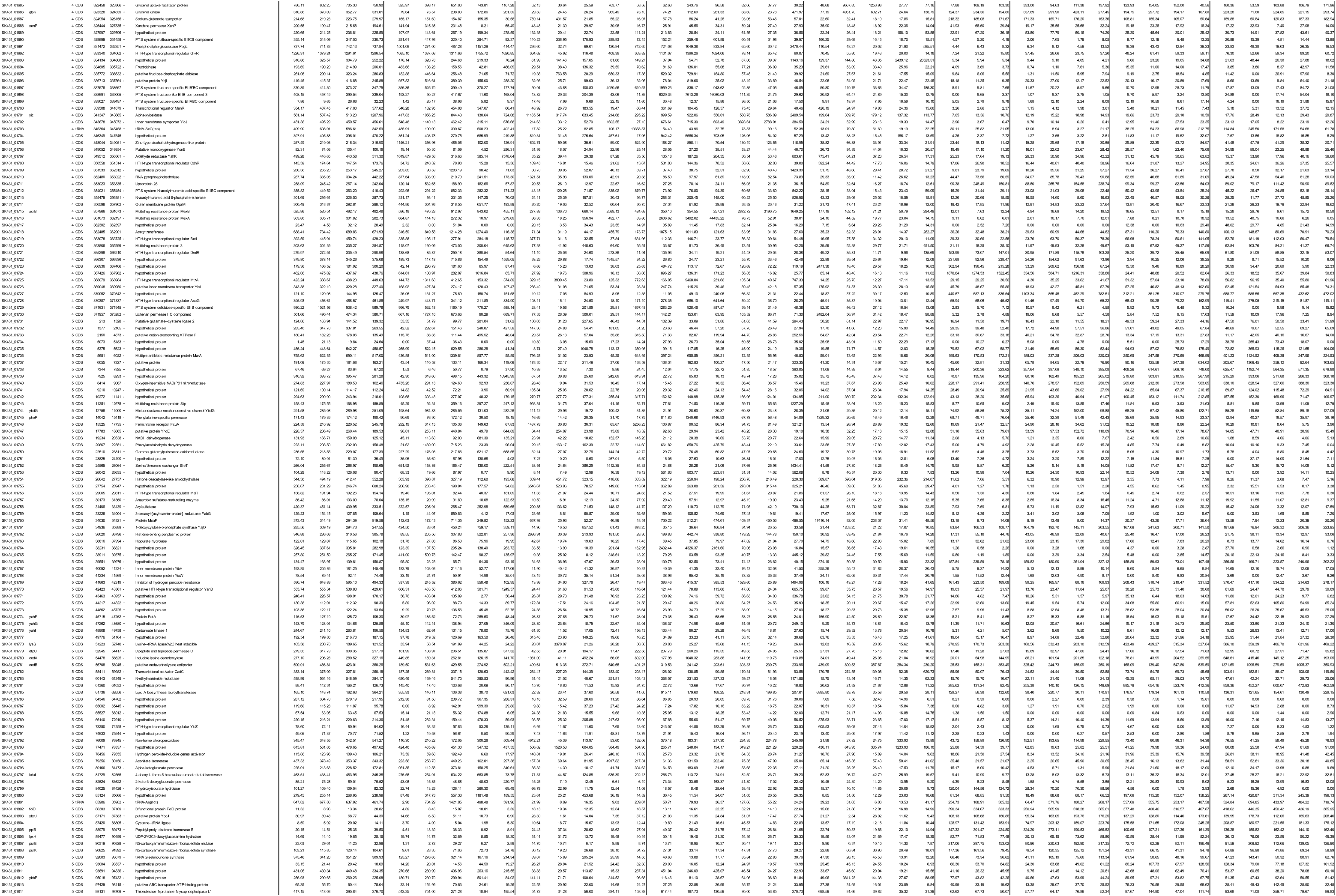
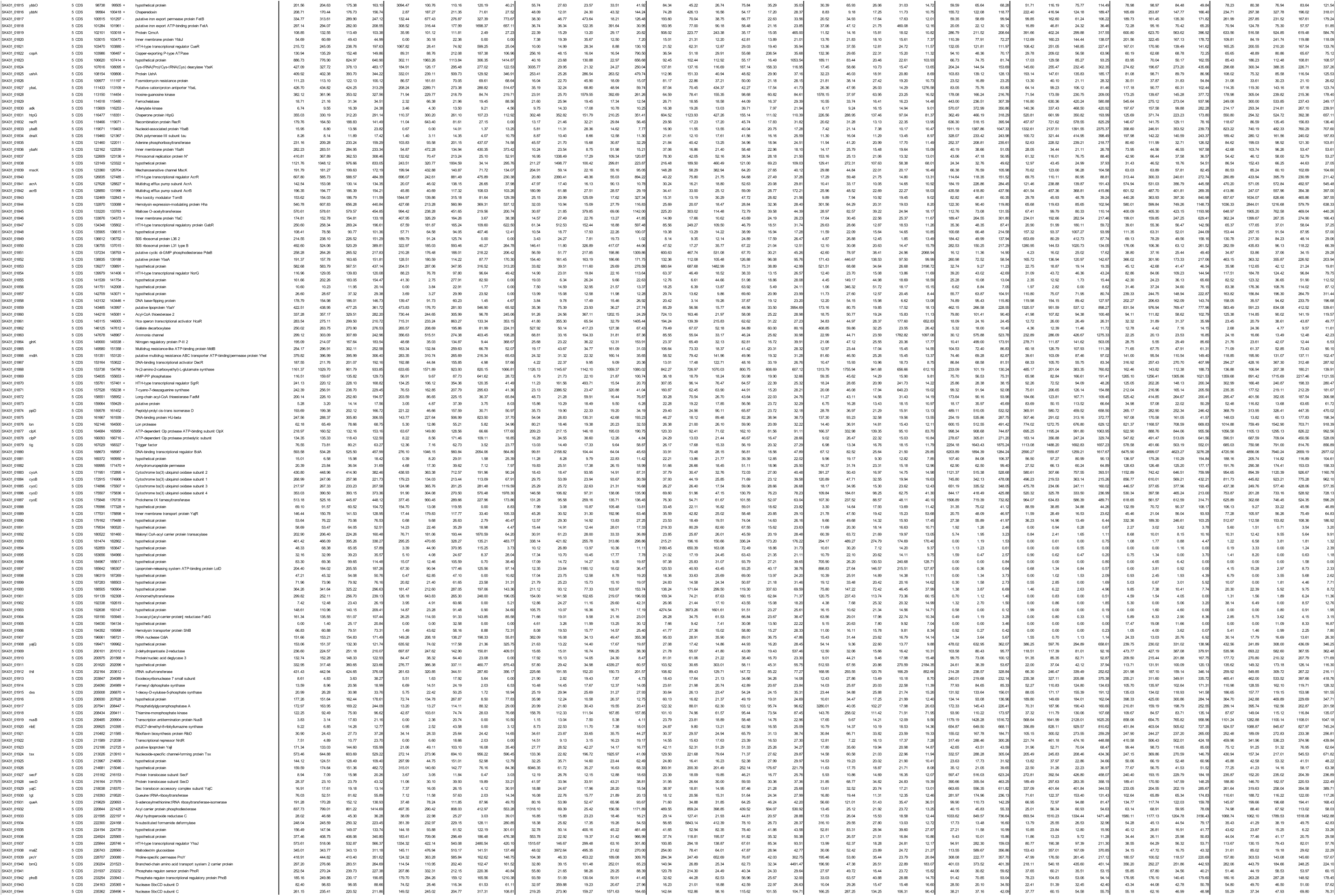
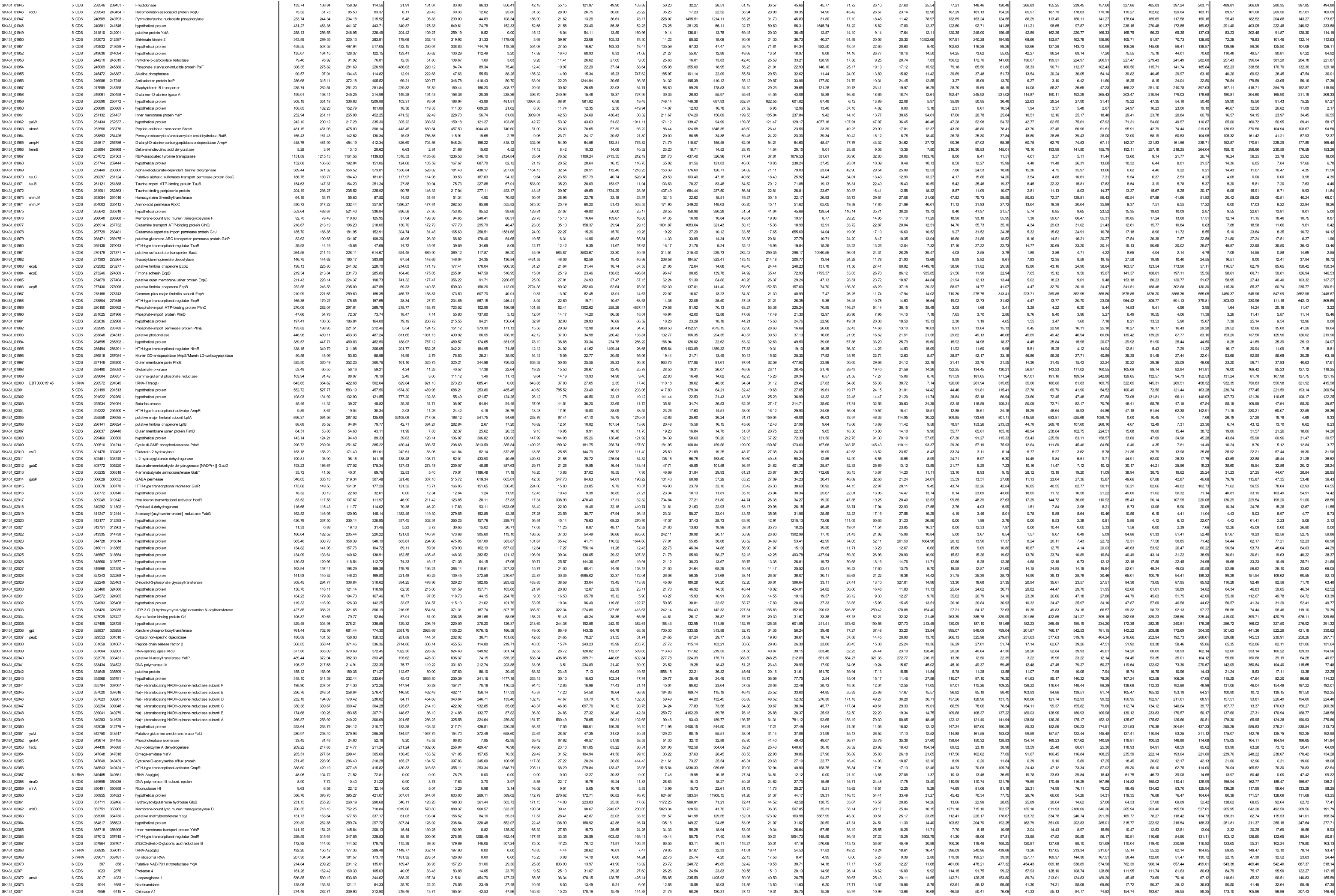
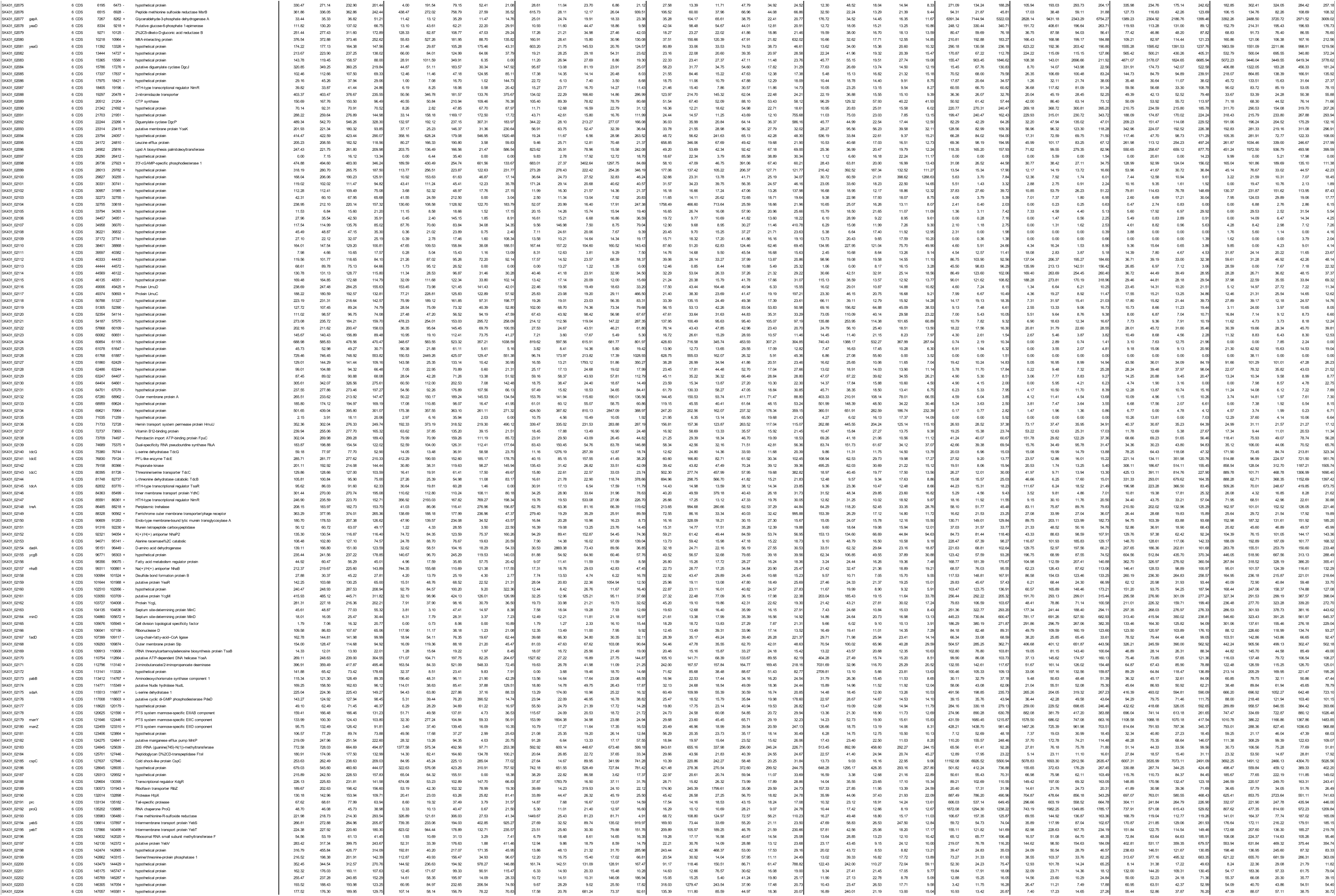
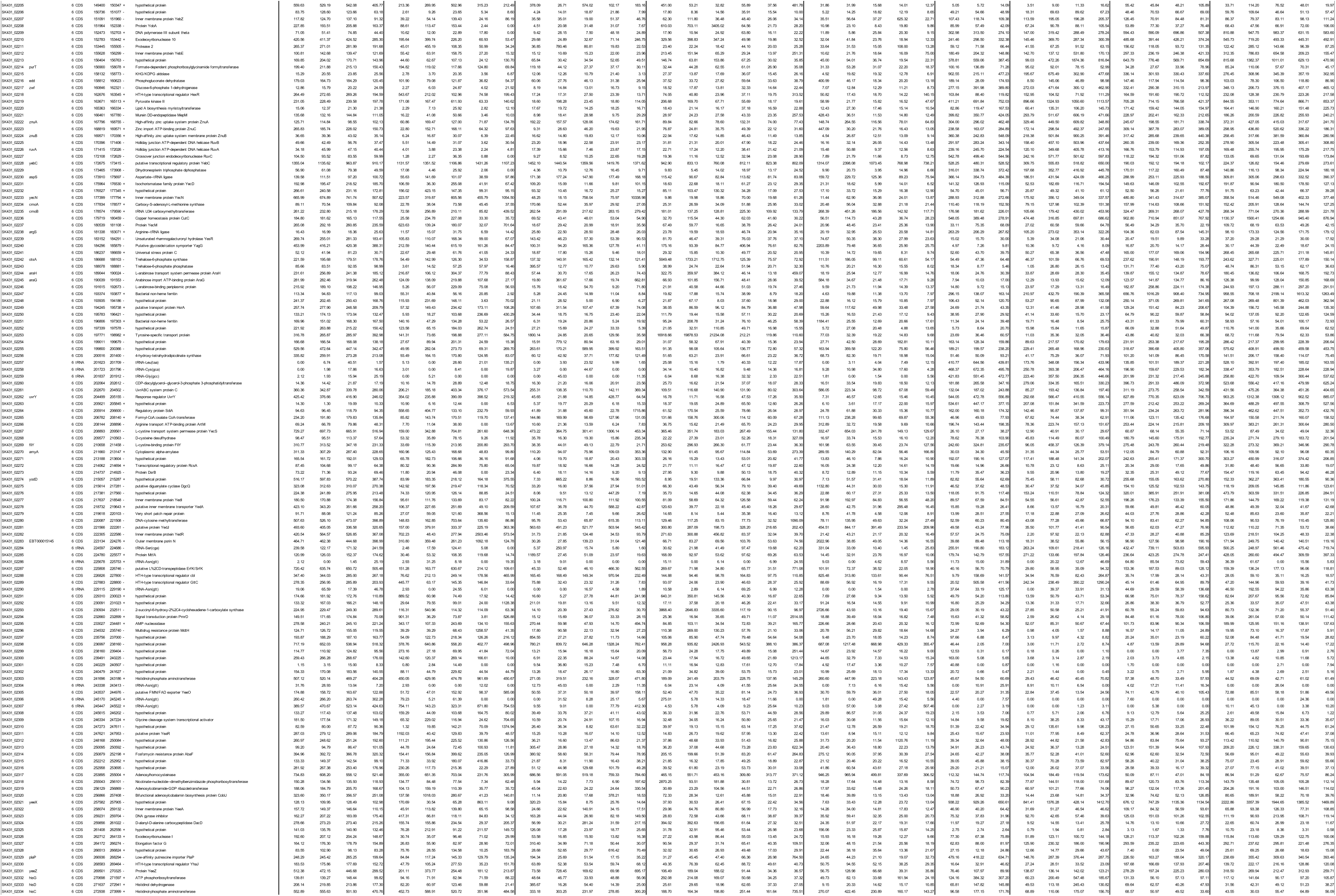
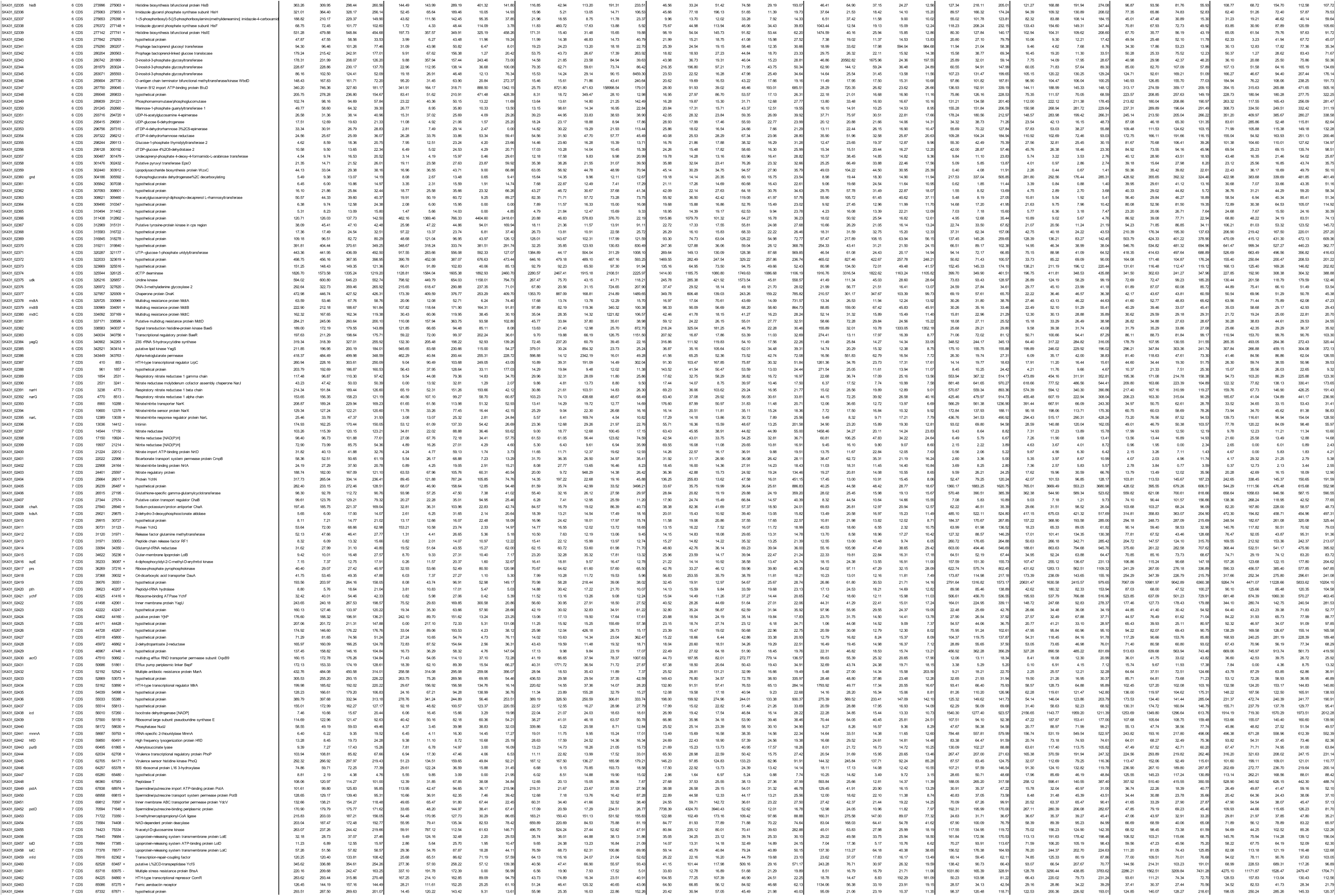
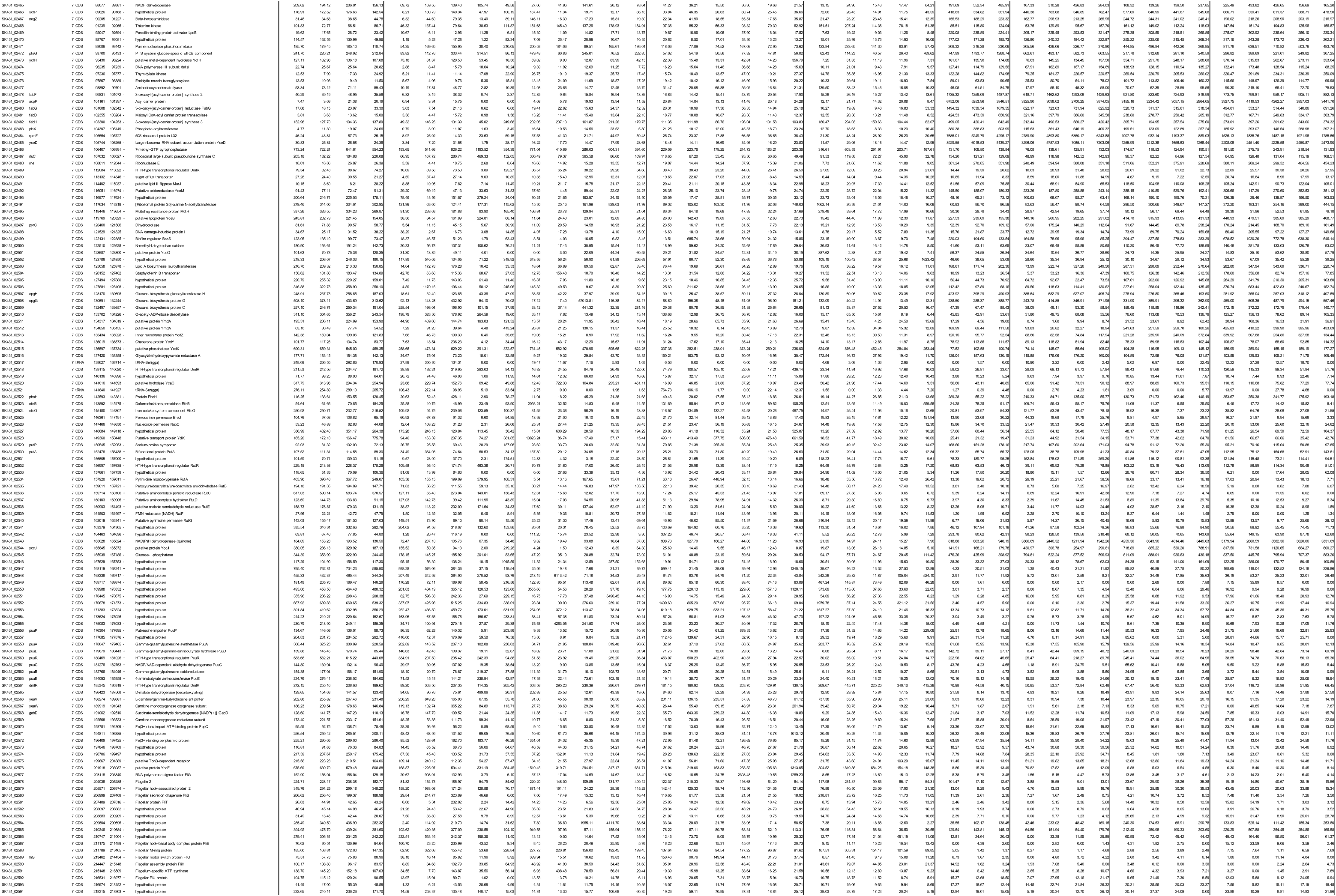
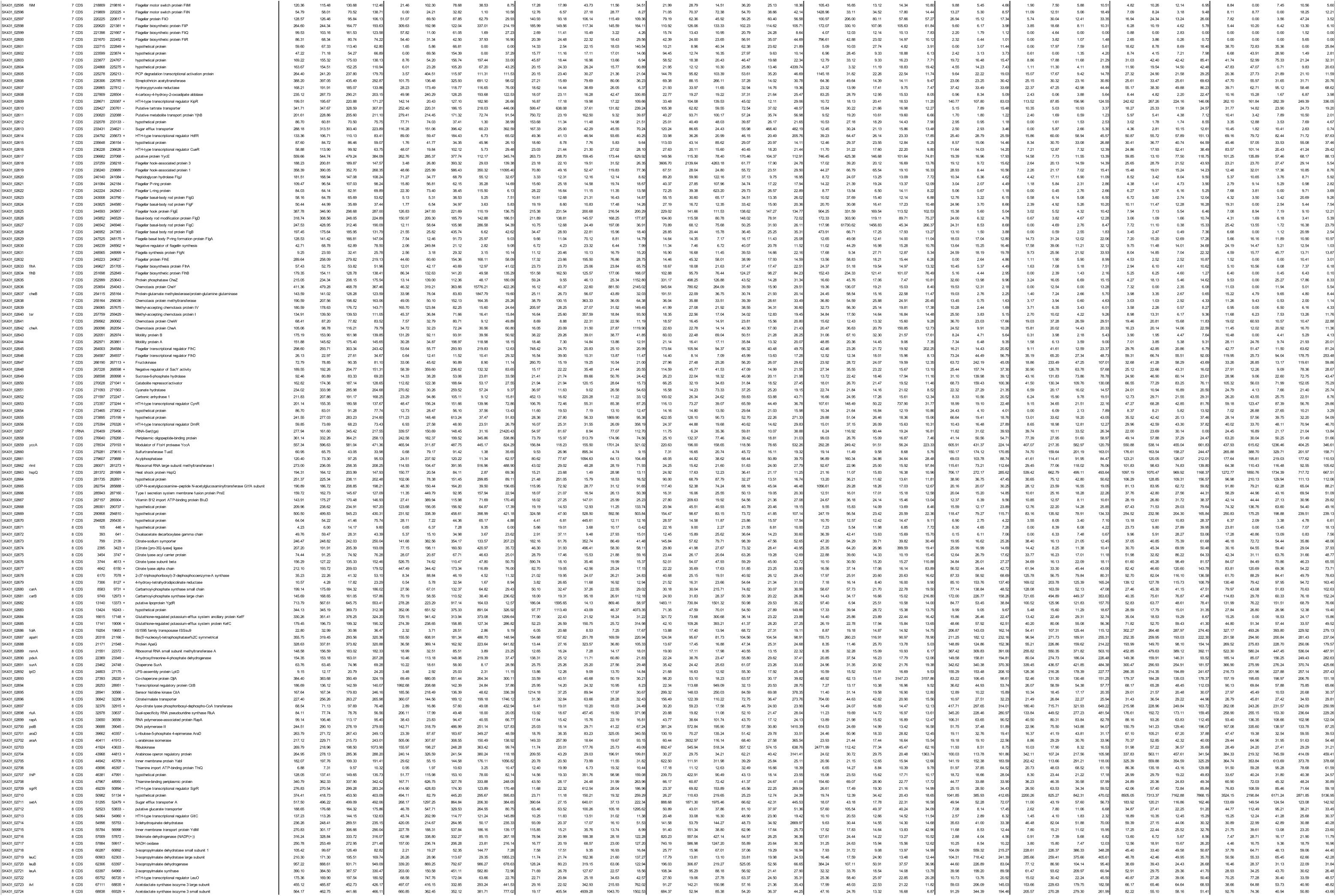
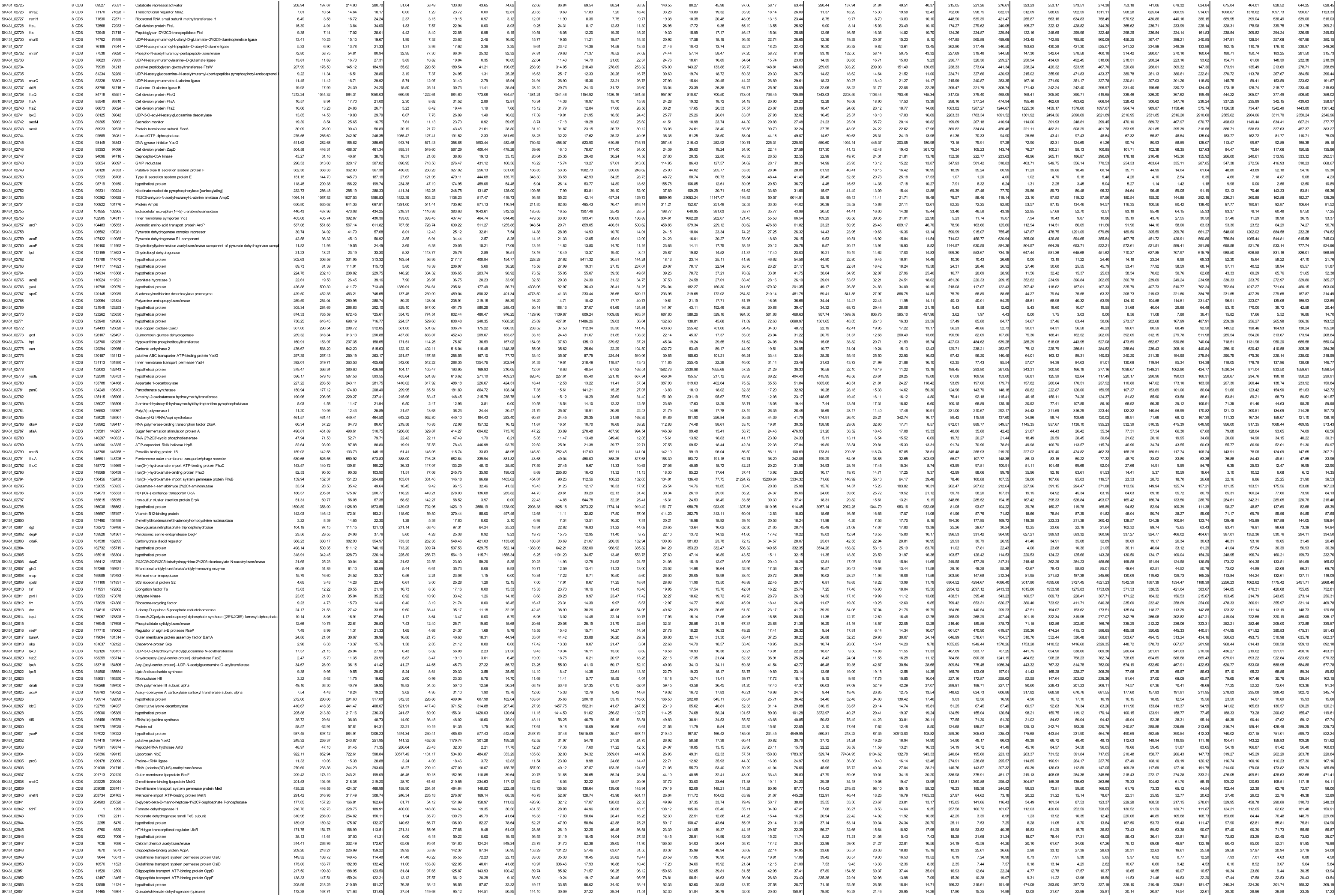
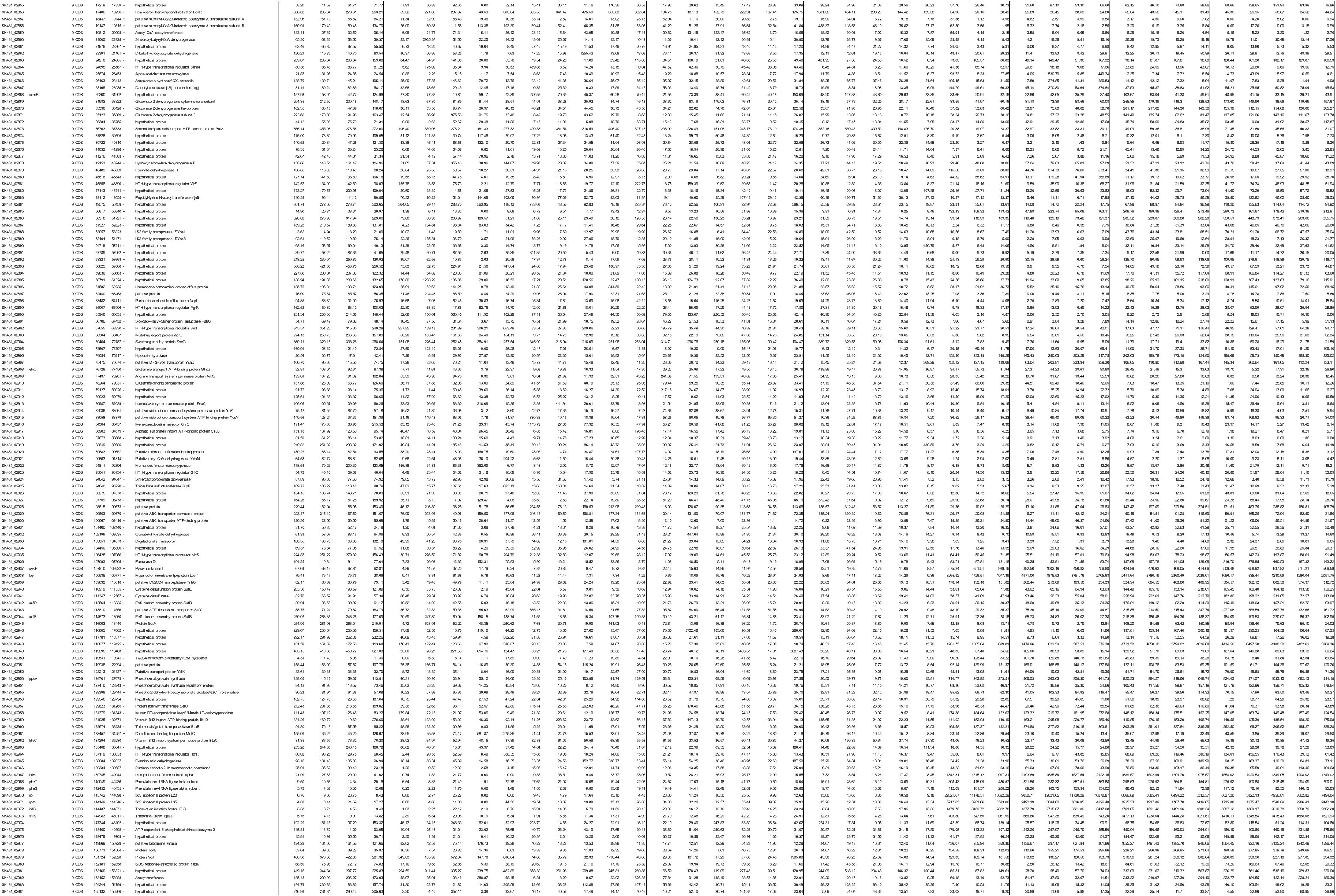
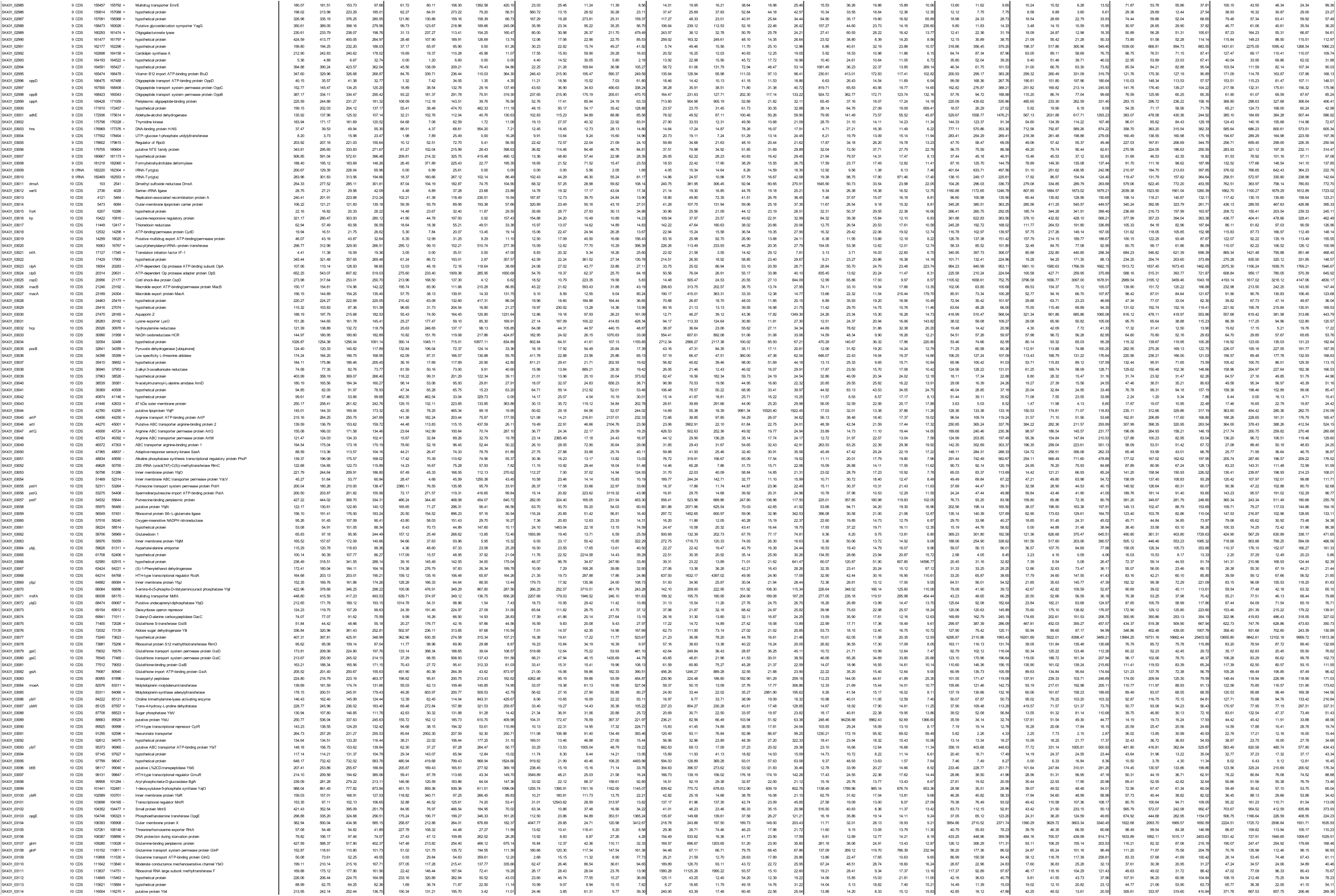
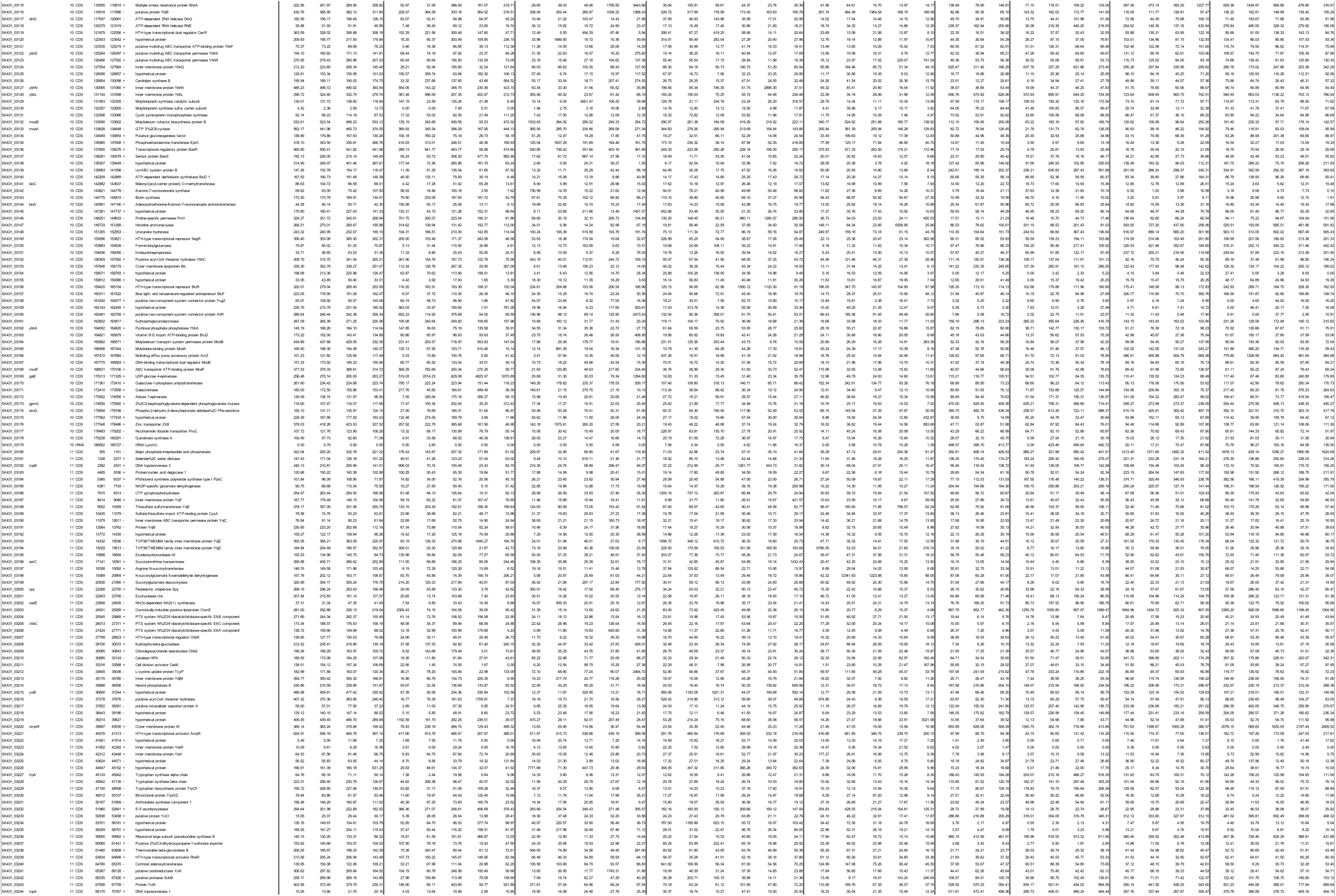
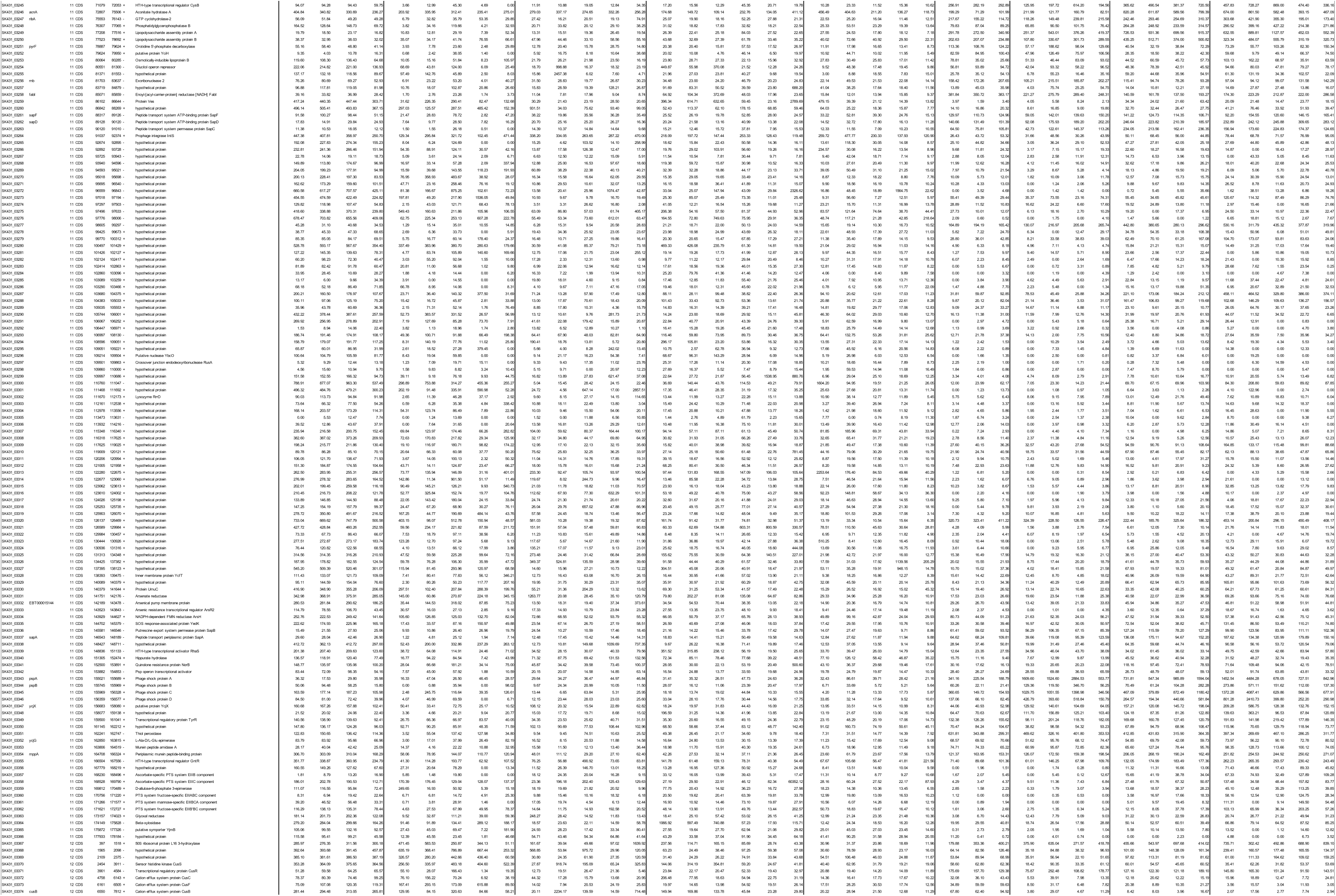
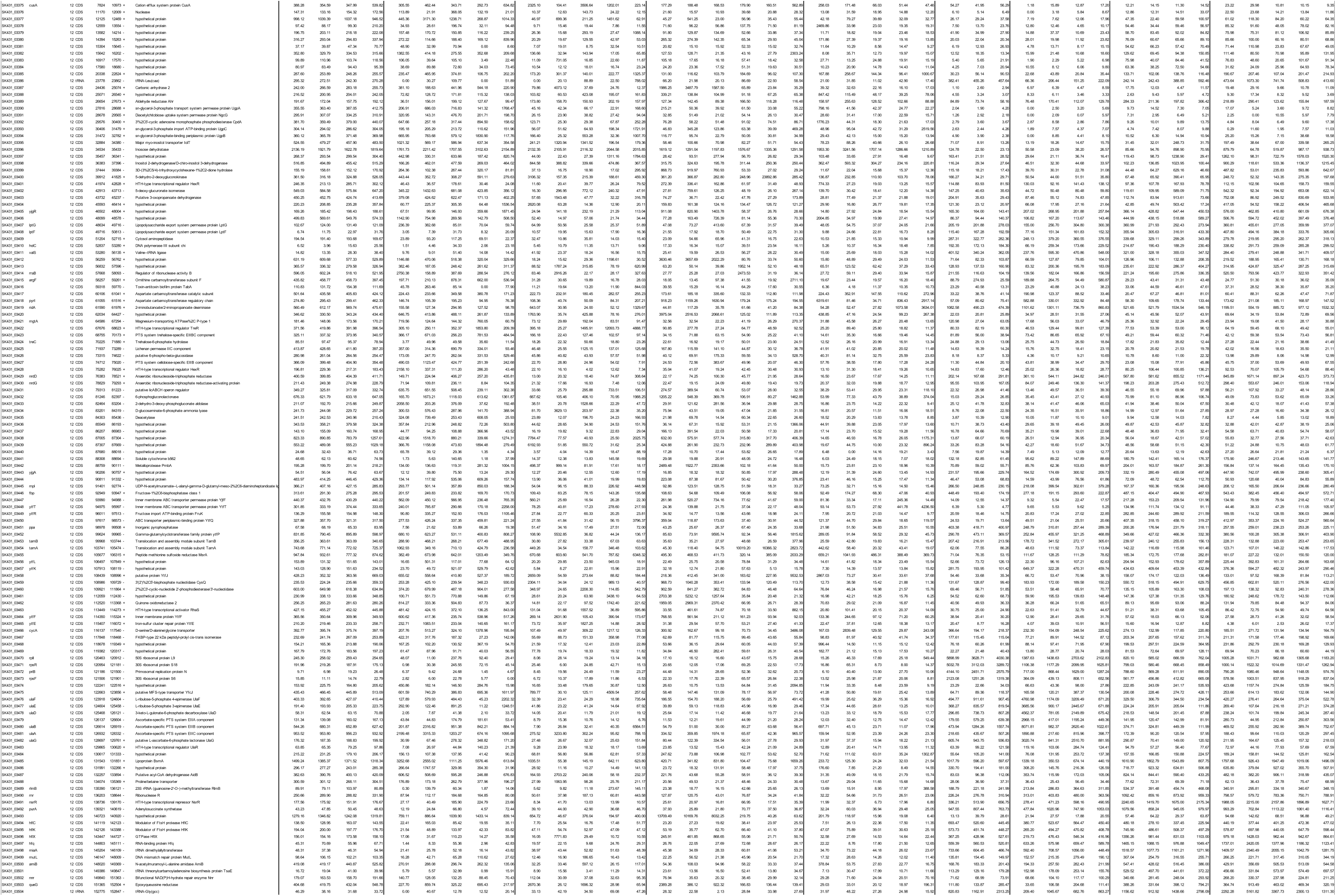
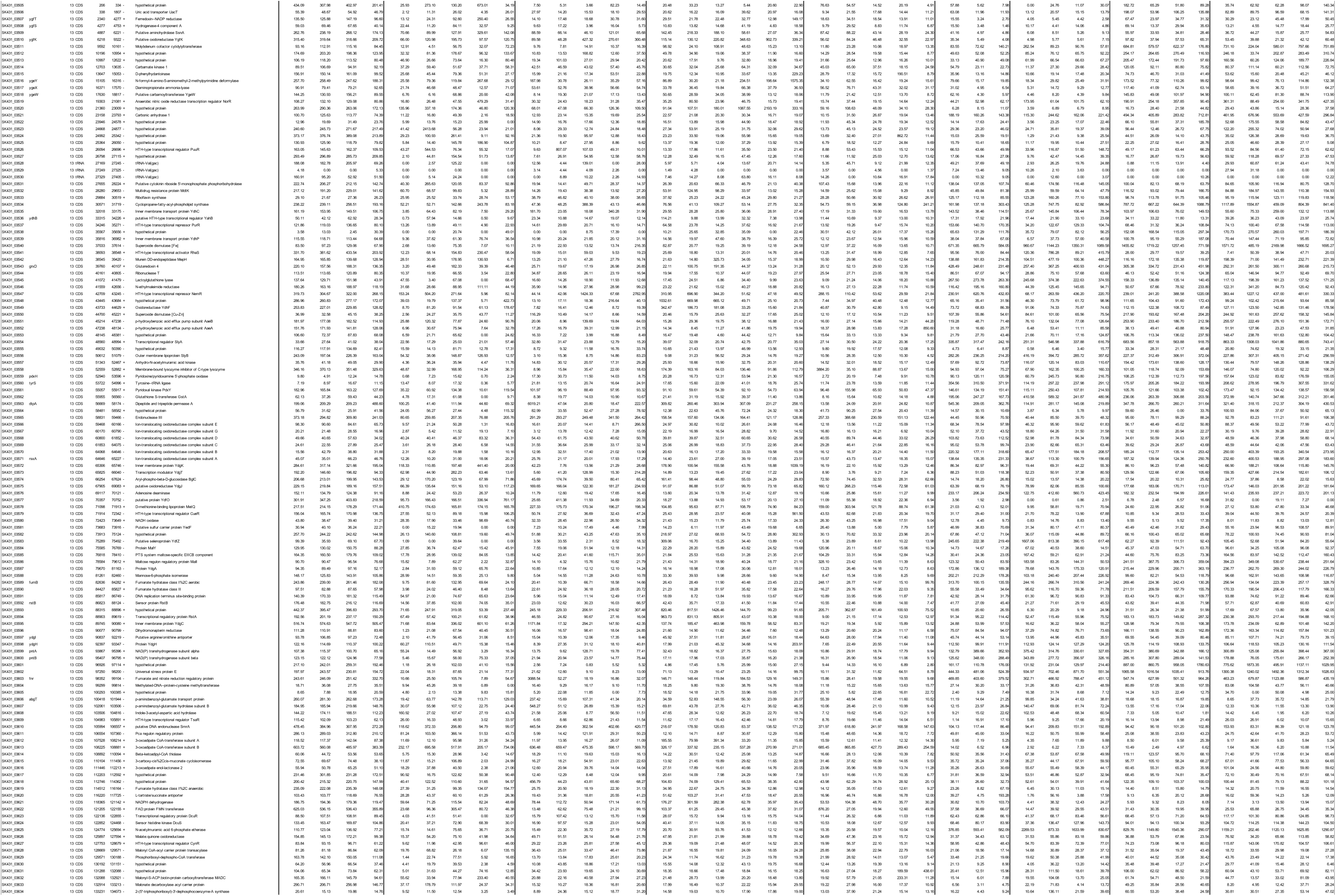
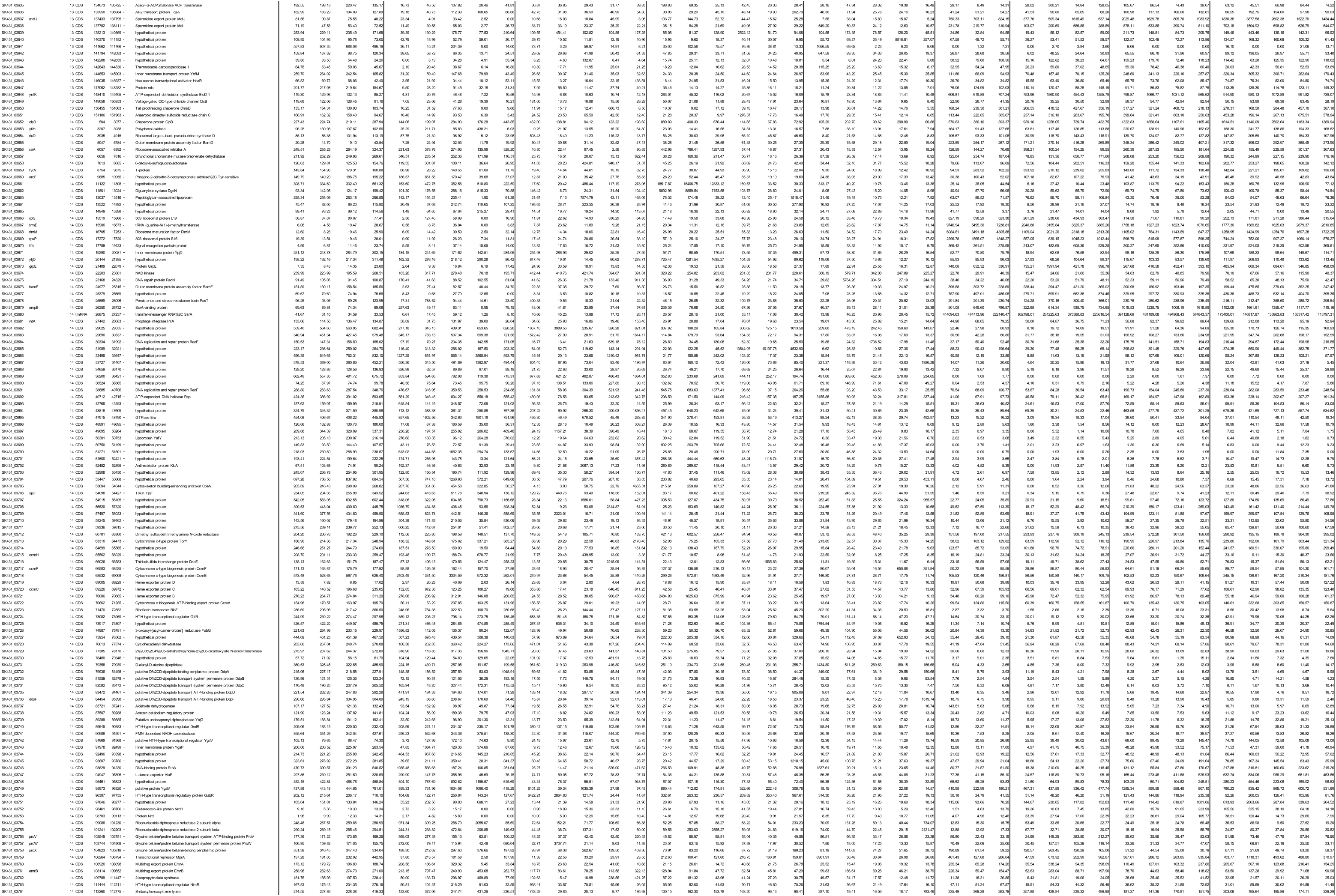
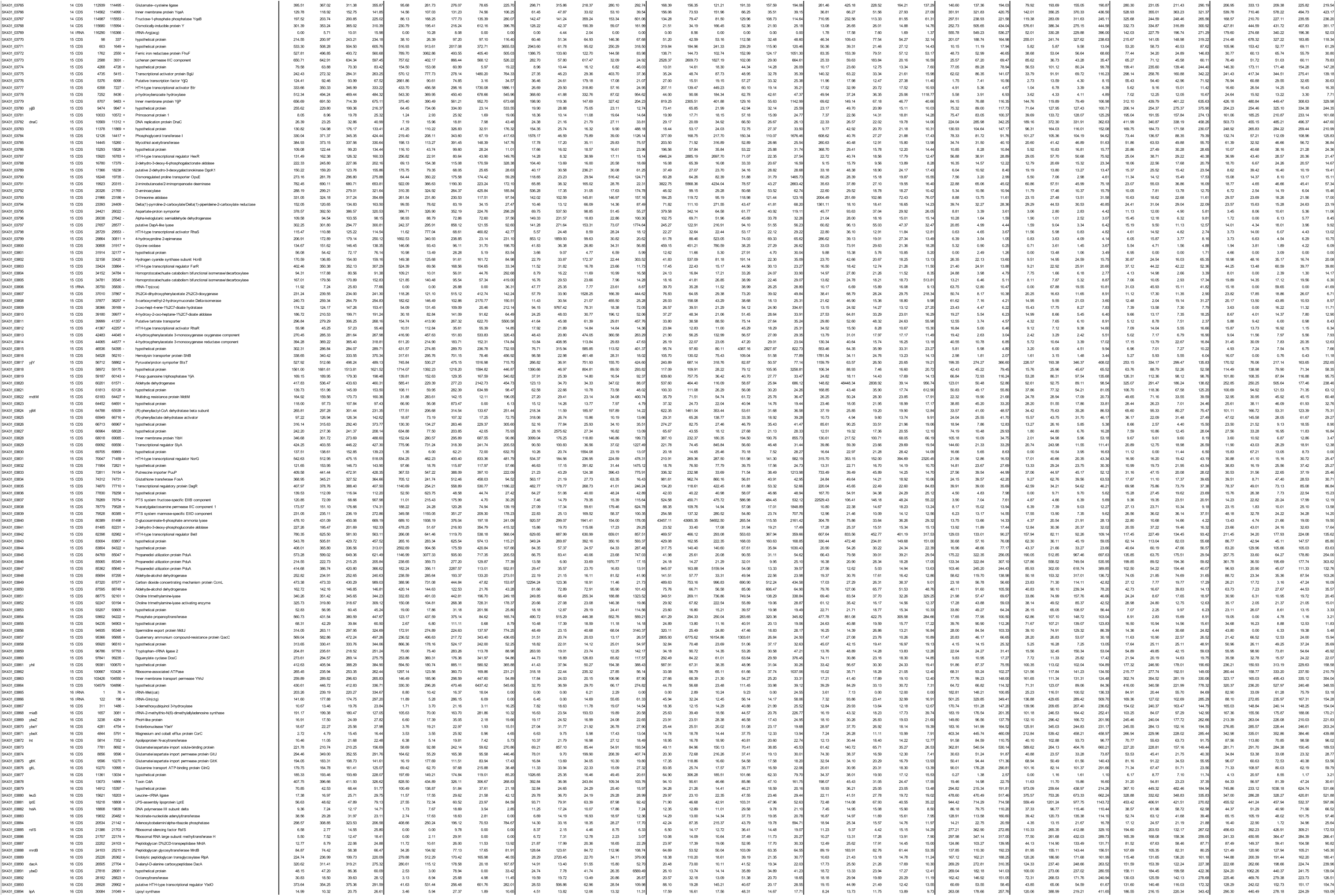
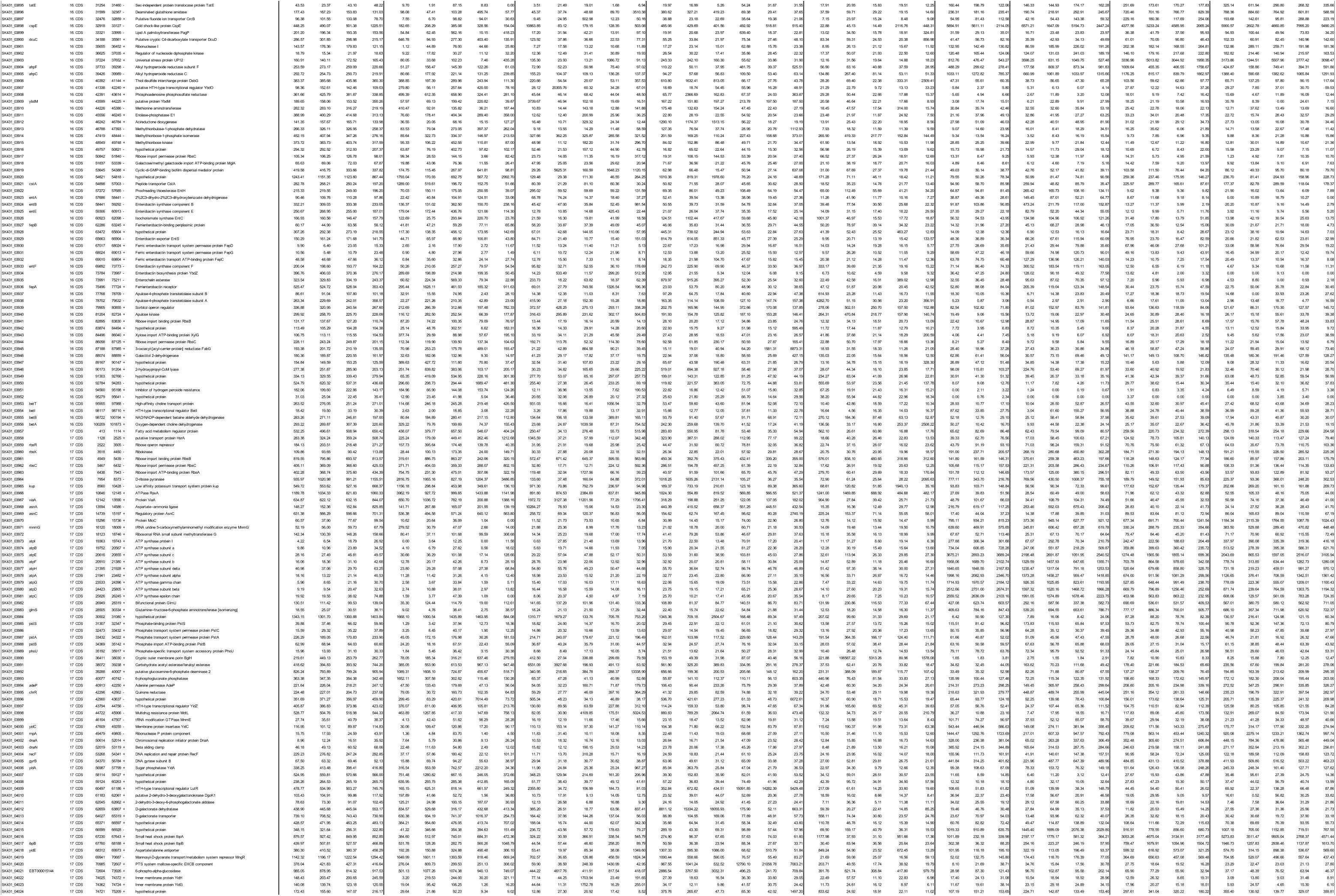
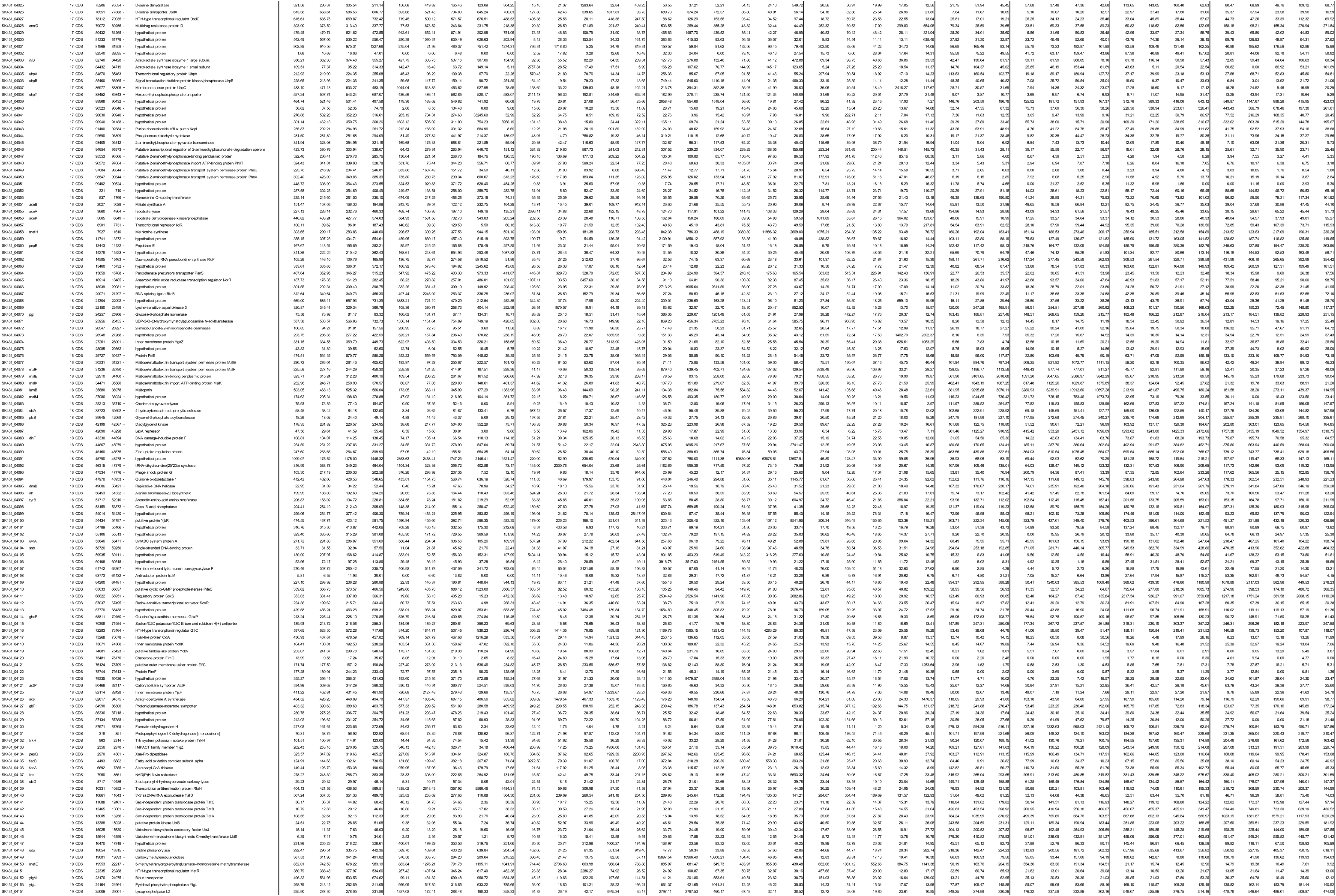
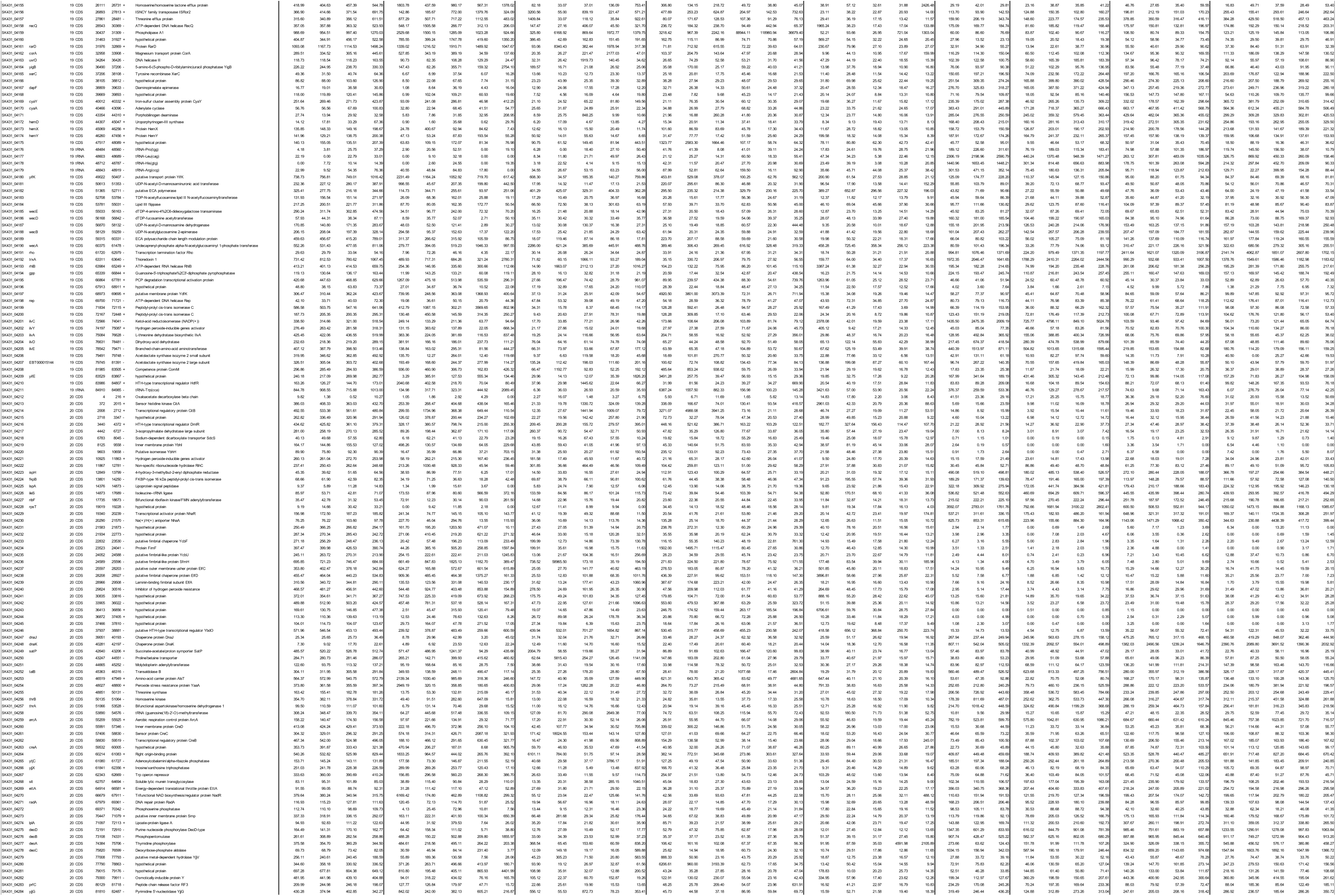
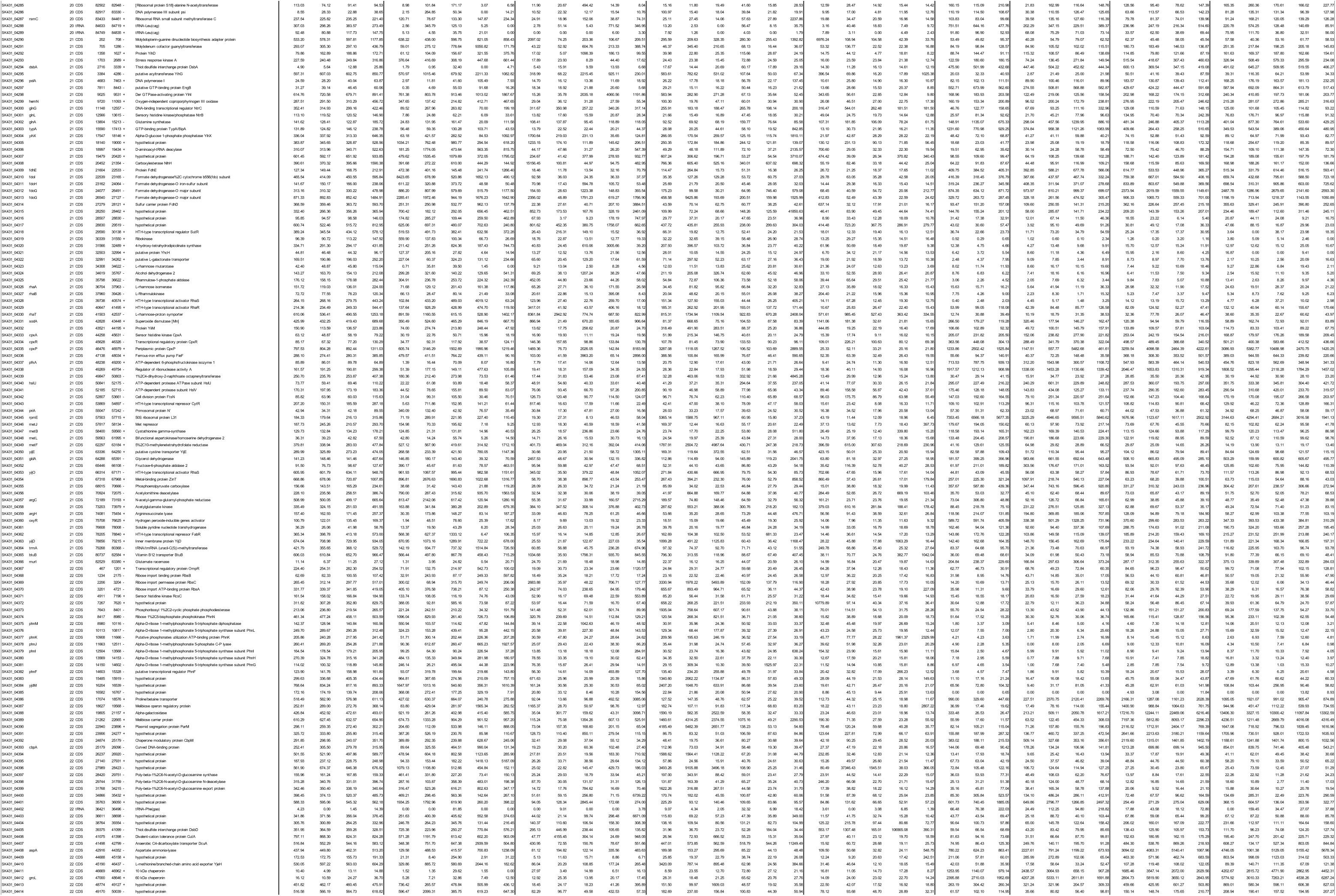
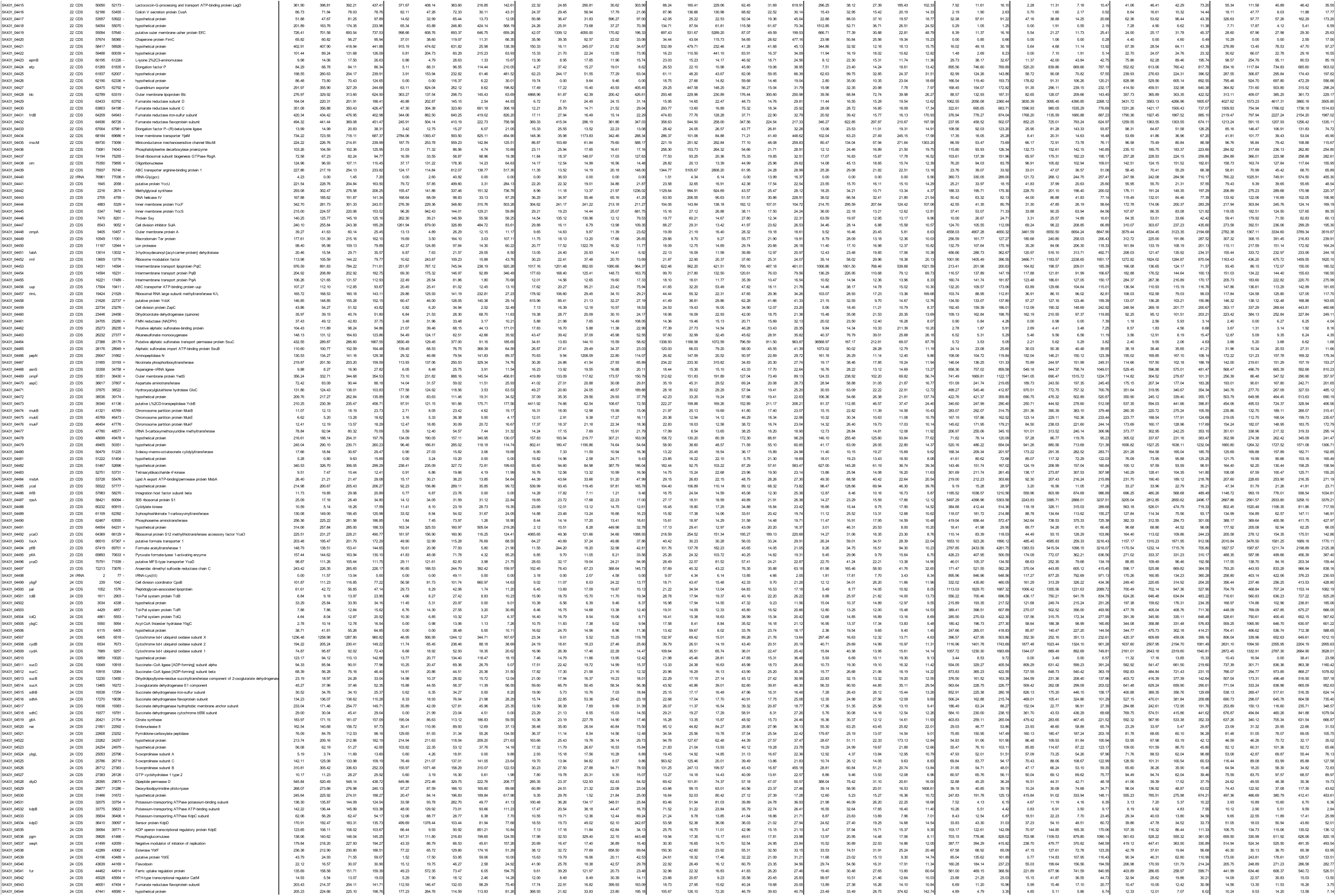
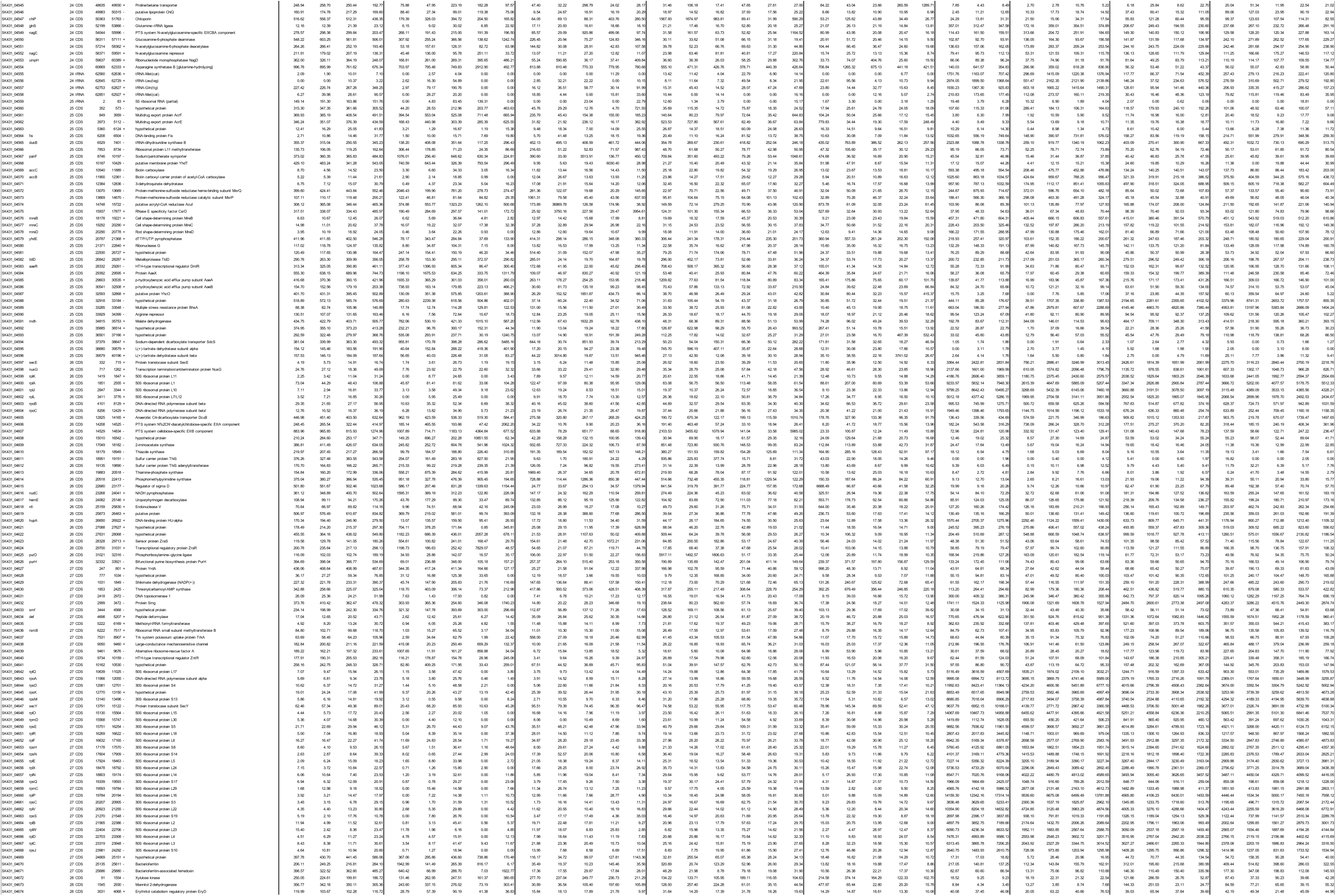
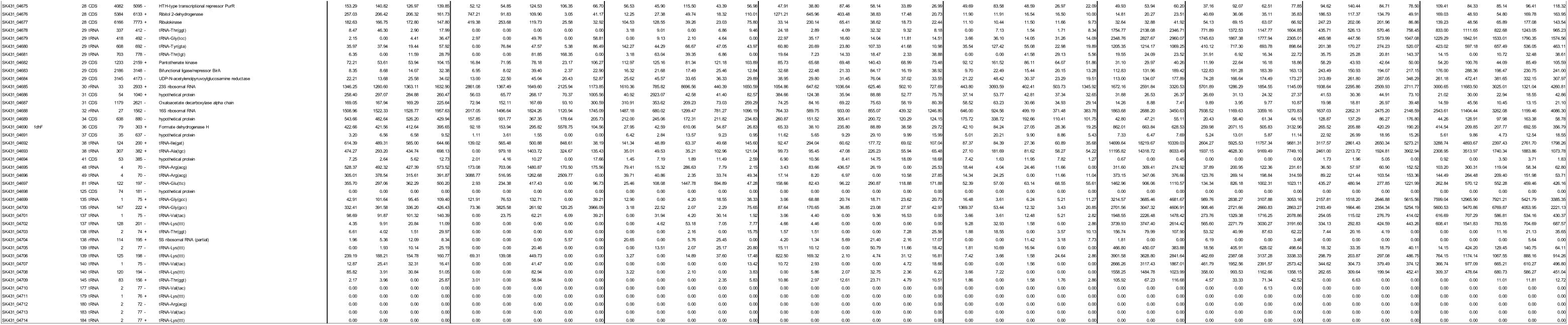
Raw data of Tn-Seq and RNA-seq.

